# Integrated hiPSC-based liver and heart microphysiological systems predict unsafe drug-drug interaction

**DOI:** 10.1101/2020.05.24.112771

**Authors:** Felipe T. Lee-Montiel, Alexander Laemmle, Laure Dumont, Caleb S. Lee, Nathaniel Huebsch, Verena Charwat, Hideaki Okochi, Matthew J. Hancock, Brian Siemons, Steven C. Boggess, Ishan Goswami, Evan W. Miller, Holger Willenbring, Kevin Healy

## Abstract

Microphysiological systems (MPSs) mimicking human organ function *in vitro* are an emerging alternative to conventional cell culture and animal models for drug development. Human induced pluripotent stem cells (hiPSCs) have the potential to capture the diversity of human genetics and provide an unlimited supply of cells. Combining hiPSCs with microfluidics technology in MPSs offers new perspectives for drug development. Here, the integration of a newly developed liver MPS with a cardiac MPS—both built with the same hiPSC line—to study drug-drug interaction (DDI) is reported. As a prominent example of clinically relevant DDI, the interaction of the arrhythmogenic gastroprokinetic cisapride with the fungicide ketoconazole was investigated. As seen in patients, metabolic conversion of cisapride to non-arrhythmogenic norcisapride in the liver MPS by the cytochrome P450 enzyme CYP3A4 was inhibited by ketoconazole, leading to arrhythmia in the cardiac MPS. These results establish functional integration of isogenic hiPSC-based liver and cardiac MPSs, which allows screening for DDI, and thus drug efficacy and toxicity, in the same genetic background.

## Introduction

The drug development pipeline generally starts with a broad collection of candidate compounds that are narrowed down by traditional *in vitro* drug screening tools including biochemical analysis, cell-based assays, e.g., using immortalized cell lines, computational modeling, and then testing in animal models. The successful compounds are then tested in human preclinical and clinical trials. The costs for pharmaceutical companies to develop a new drug from initial compound screening to market is estimated at more than 2.5 billion USD^1, 2^, and can take ~10-15 years^3, 4^. One of the more costly and inefficient steps in this process is the clinical trial phase^5^. Too often compounds pass the go/no-go risk decisions based on animal model data, but fail later in clinical trials due to species-specific differences in physiology. The worst-case scenario occurs when drugs make it through the full screening pipeline and clinical trials to market, only to be recalled due to unforeseen side effects in some patients, a prominent example being the arrhythmogenic effect of the gastroprokinetic cisapride^6^. In addition to inability of animal models to accurately mimic human physiology, such failures are due to lack of diversity in the human genetic backgrounds tested in single cell lines, and lack of diversity in phase I and II clinical trials.

Drug-induced liver injury (DILI) and cardiotoxicity are the leading causes of preclinical and clinical withdrawals of marketed pharmaceuticals due to safety issues^7, 8^, and manifest as serious complications, including hepatitis and liver cell death, potentially leading to liver failure, and cardiac arrhythmia^9, 10^. The human liver is especially significant for toxicity as it is the first stop for most compounds after they enter the body for initial drug metabolism before subsequent uptake by other organs^11^. Animal models perform variably in representing human diseases and predicting toxicity to humans due to inter-species differences in drug absorption, distribution, metabolism, excretion, and/or toxicity (ADME/Tox)^12, 13, 14^. Accordingly, the risk of human DILI is not well predicted by animal models. Additionally, animal studies can only partially recapitulate patient-specific factors such as genetics, age, environment, and concomitant diseases^15^.

The limitation of animal models to predict DILI has led to the development of *in vitro* liver models; however, current versions have limitations, most notably due to the complexity of the liver, which performs more than 500 functions, including metabolic, synthetic, immunologic, and detoxification processes^16^. The most commonly used models of liver toxicity involve two-dimensional (2D) monolayer cultures of primary human hepatocytes (pHeps). They currently represent the gold standard for drug screening, toxicology studies, cell-based therapies, and *in vitro* disease modeling. Indeed, they recapitulate most liver functions and represent diverse human genetic disease backgrounds that are not captured by single human cancer cell lines. pHeps also bypass ethical concerns regarding refined and reduced use of animal experiments. However, there are still substantial problems using pHeps. One of the limitations is the difficulty to perform long-term cultures (≥7 days) due to the propensity of pHeps to dedifferentiate *in vitro* within a few days, causing them to lose hepatocyte-specific functions^17, 18^ and no longer represent accurately the biology of the liver. Recent advances in long-term cultures of pHeps might overcome these limitations^19^ but not the shortage of donor livers^20, 21^, resulting in high cost and considerable genetic and functional variation of commercial pHeps^22^. These variations alone make interlaboratory comparisons nearly impossible. Taken together, conventional 2D monolayer cultures using pHeps are often too simplified, ultimately failing to adequately predict DILI, putting human lives at risk and wasting valuable resources for pharmaceutical companies during clinical drug development.

Human induced pluripotent stem cell (hiPSC)-derived hepatocytes (hiPSC-Heps) have the potential to overcome the limitations of pHeps and human cell lines. hiPSCs are pluripotent cells generated by reprogramming of mature somatic cells, e.g., adult human fibroblasts, by overexpression of the Yamanaka factors^23, 24^. hiPSCs can provide a virtually inexhaustible source of cells from different genotypes, including healthy or disease-specific cells^25, 26^. By providing a renewable source of cells that maintain the genotype of the donor, hiPSCs reduce cell source variability and facilitate an unlimited number of trials of patient-specific drug screening at the early-stage of the development pipeline^27, 28, 29, 30^. However, like other protocols for directed differentiation of hiPSCs, current differentiation protocols for hiPSC-Heps produce immature cells that lack certain functions of pHeps.

Although pHeps and hiPSC-Heps are widely used in 2D monolayer culture for initial assessment of drug metabolism, these monocultures do not provide three-dimensional (3D) organization, non-parenchymal cells, structural cell-cell interactions, nor their associated paracrine interactions^31^. Advances in 2D coculture (e.g., of hiPSC-Heps and fibroblasts) exploiting micropatterned architecture have shown promising results predicting DILI^32^, but improvement is necessary to achieve accurate prediction of the human liver. Compared to 2D, bioinspired 3D *in vitro* systems provide more structural organization, which is beneficial to maintaining cell differentiation and maturity^33,34^. For example, 3D culture can reverse or prevent dedifferentiation of pHeps^35, 36^ and promote metabolic enzyme functionality comparable to *in vivo* levels for adult rat hepatocytes^14, 37, 38^. Thus, the combination of hiPSCs and 3D culture offers great potential to improve the efficiency and efficacy of the drug development pipeline^39, 40, 41^ by allowing to study pHeps in a more biologically relevant setting. However, only few published studies used hiPSC-Heps to build 3D liver microphysiological systems (MPSs)^31, 42^. Furthermore, a challenge not addressed by previous research is linking the liver with other organs to study inter-organ crosstalk, which is important for studying DDI.

Here, we present the development of a hiPSC-based *in vitro* liver model on a microfabricated platform, i.e., MPS (also known as ‘organ-on-a-chip’ or ‘tissue chip’), fluidically integrated with a previously developed hiPSC-based cardiac MPS platform^4, 43, 44^. Using hiPSCs allowed us to produce different 3D tissues—liver and heart—from the same donor and thus with identical genetic background. In addition, we modified our recently reported protocol^45^ to produce hiPSC-Heps suitable for studies of drug metabolism, including sufficient cytochrome P450 (CYP) 3A4 activity and long-term function in the MPS. As an example of clinically relevant DDI, we focused on cisapride, a gastroprokinetic intended to treat the symptoms of heartburn in adults^46^. This drug causes prolongation of the QT interval, ventricular arrhythmias, and torsade de pointes^47^ in patients taking medications that interfere with cisapride metabolism or prolong the QT interval. In 2000, cisapride was removed from the market as a result of 23 deaths^48, 49^, and 341 incidences of torsade de pointes. To fluidically combine the liver and cardiac MPSs, we developed a common medium for survival and function of each tissue. By correctly predicting CYP3A4-dependent DDI between cisapride and the antifungal ketoconazole, we show that MPS technologies are effective into predicting drug efficacy and toxicity across multiple organs, and have potential as next-generation high-content drug development tools.

## Results

### Directed differentiation of hiPSC-Heps

Directed differentiation of hiPSCs into hiPSC-Heps was performed in 2D cell culture using an optimized protocol (**Fig. 1a**; see Material and Methods for details). Brightfield microscopy showed characteristic morphological changes occurring during the differentiation process (**Fig. 1b**) from hiPSCs (day 0) to definitive endoderm (day 8), liver progenitors (day 13), immature (day 18) and mature hepatocytes (day 23). Immunofluorescence showed lack of the pluripotency marker OCT3/4 past the hiPSC stage (**Fig. 1b**); hepatic nuclear factor 4 alpha (HNF4A) started to be expressed at day 13 when cells committed to hepatic fate; alpha-fetoprotein (AFP) stained strongly positive in immature hepatocytes at day 18; albumin (ALB) was only detectable at the final stage of differentiation (day 23). At this stage, 88.4% of cells expressed albumin and 32.7% were positive for asialoglycoprotein receptor 1 (ASGR1), a marker of mature hepatocytes, in flow cytometry (**Fig. 1c** and **Supplementary Fig. 1**).

**Fig 1.**
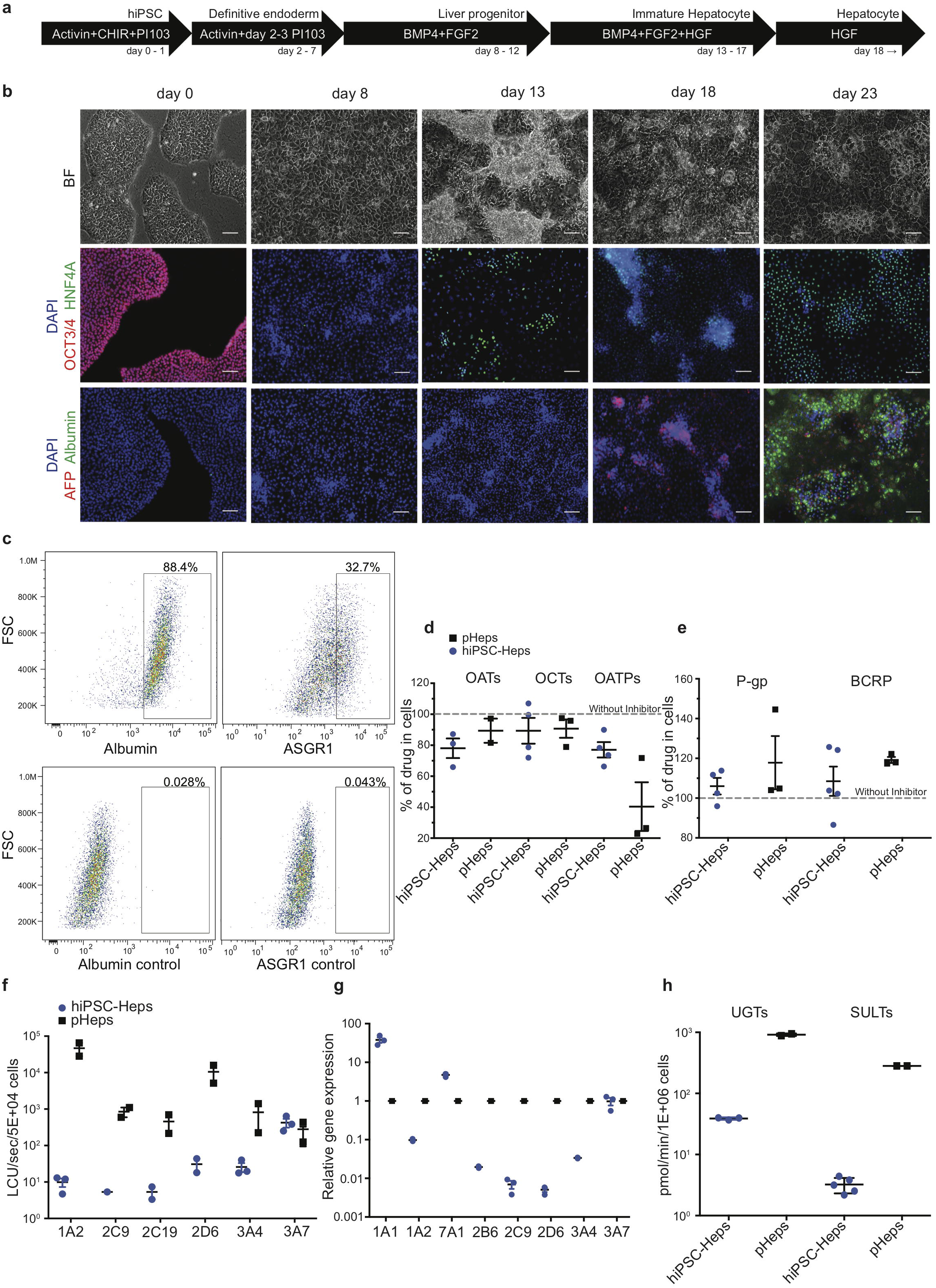
Differentiation and characterization of hiPSC-Heps. a) Schematic of the hiPSC-Heps differentiation protocol. Arrows show cells at different stages progressing from hiPSCs (day 0) to mature hepatocytes (day 23). Growth factors and small molecules (PI-103 and CHIR) are listed inside the arrows. b) Brightfield and immunofluorescent images matching the differentiation stages in Fig. 1a. Nuclei were stained with DAPI. Scale bar, 100 μm. c) Flow cytometry shows percentage of hiPSC-Heps expressing albumin and ASGR1 at day 23. d) Activity analysis of uptake transporters. Transporter activity is presented as the decrease of drug concentration in the cells in the presence of the inhibitor (% of drug in cells). Data are presented as the mean +/− standard deviation from three to four (hiPSC-Heps) or two to three (pHeps) independent experiments. e) Activity analysis of efflux transporters. Results obtained with inhibitors were set to 100% to display transporter activity as increase of drug concentration in the cells. Data are presented as the mean +/− standard deviation from four to five (hiPSC-Heps) and three (pHeps) independent experiments. f) Activity analysis of CYPs. Data are presented as the mean +/− standard error of the mean from three (hiPSC-Heps) or two (pHeps) independent experiments. g) Quantitative reverse transcription PCR analysis of CYP genes. h) Activity analysis of conjugating enzymes. Data are presented as the mean +/− standard deviation from two (UGTs, SULTs; pHeps), three (UGTs; hiPSC-Heps) or five (SULTs; hiPSC-Heps) independent experiments.

### Characterization of drug metabolism in hiPSC-Heps

A prerequisite for drug metabolism studies is sufficient cellular uptake of the drug as well as efflux of the metabolites. Therefore, activity of three uptake (**Fig. 1d**) and two efflux (**Fig. 1e**) transporters was measured by high performance liquid chromatography coupled with tandem mass spectrometry (LC-MS/MS) using drug/inhibitor combinations. For the uptake transporter studies, hiPSC-Heps were exposed to a specific drug internalized by the transporter, with or without a specific inhibitor of the transporter—activity of organic anion transporters (OATs) was tested with acyclovir/probenecid, organic cation transporters (OCTs) with metformin/decynium-22 and organic anion transporting polypeptides (OATPs) with rosuvastatin/rifamycin-SV. Metabolism-qualified pHeps were used as controls. All three phase 0 uptake transporters were active in hiPSC-Heps and pHeps, including responsiveness to specific inhibitors (**Fig. 1d**). Next, the activity of two phase III efflux transporters was analyzed using substrate/inhibitor challenge. The P-glycoprotein (P-gp) was tested with the substrate digoxin and the breast cancer resistance protein (BCRP) with mitoxantrone. Both efflux transporters were active in hiPSC-Heps and pHep, including responsiveness to the P-gp/BCRP inhibitor GG918 (**Fig. 1e**). To further assess whether our hiPSC-Heps allowed studies of phase I drug metabolism, we measured the activity of key hepatic CYP enzymes (**Fig. 1f**). Based on a luminescence assay, CYP3A4 was one of the most active drug-metabolizing CYPs in hiPSC-Heps and its activity was in the range of 3 to 5% of pHeps (**Fig. 1f**). Activities of other CYPs were 2-3 orders of magnitude lower in hiPSC-Heps than in pHeps, except for CYP3A7 (**Fig. 1f**). High activity of “fetal” CYP3A7 was confirmed by gene expression analysis and, together with CYP1A1/CYP1A2 ratio, indicated that the majority of our hiPSC-Heps resembled fetal or immature hepatocytes (**Fig. 1g**). In accord, the activities of the phase II conjugating enzymes UDP-glucuronosyltransferases (UGTs) and sulfotransferases (SULTs) were in the range of 1 to 10% of pHeps, as shown by LC-MS/MS using the substrate/metabolite combination 1-naphtol/naphtol-s-glucuronide and nitrophenol/nitrophenol-sulfate, respectively (**Fig. 1h**). Viewed together, our optimized protocol produced hiPSC-Heps capable of all phases of drug metabolism, although some transporter and enzyme activities were underdeveloped. This finding reflects the current state of protocols for hiPSC-Hep differentiation and underscores the importance of in-depth characterization of these cells to ascertain suitability for the intended experiments.

### Development of the liver MPS

The liver MPS was designed to mimic a single acinus, the smallest functional unit of the liver, which is comprised of three different zones, exposing hepatocytes to different concentrations of nutrients and oxygen (**Fig. 2a**). Our chip design was a cylindrical-like structure that was cut in half and laid flat, thus creating two rectilinear chambers—one chamber for the hiPSC-Heps and one chamber for media to mimic the sinusoid. The chambers were separated by an isoporous polyethylene terephthalate (PET) membrane with 3-μm pores that acted as a diffusive barrier, mimicking the function of the liver sinusoidal endothelial cells that protect the hepatocytes from shear stress forces of the blood flow. Prodanov *et al*^50^ used a similar approach with pHeps and two additional cell lines to model contributions from endothelial cells and hepatic stellate cells. Our system used only hiPSC-Heps and relied on a different fluidic design to minimize shear stress within the hiPSC-Heps chamber, thereby reducing potential cell damage. Additionally, our liver MPS was designed to allow loading of a higher number of cells by centrifugation to improve 3D liver tissue formation and metabolic function.

**Fig 2.**
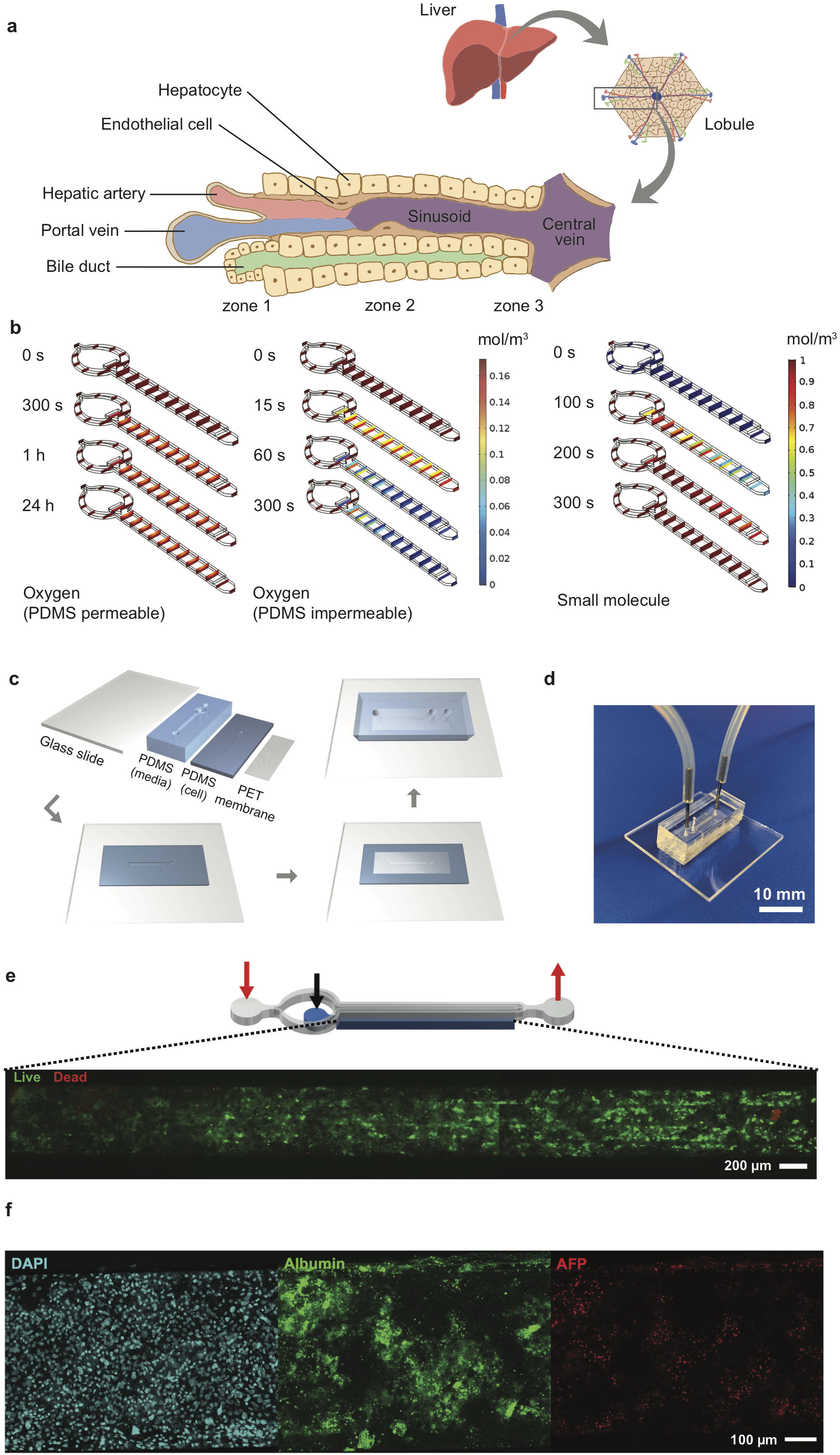
Design, characterization, fabrication and testing of liver MPS. a) Illustration showing the liver acinus as the basic building block of the liver inspiring the liver MPS design. b) COMSOL simulation results. Small concentration gradients with physiologically relevant oxygen levels are observed when cells in the cell chamber consume oxygen and oxygen diffuses from the ambient through the PDMS roof and walls (left). When oxygen is not diffused through the PDMS cells become hypoxic after 300 seconds (center). The hiPSC-Heps OCR used in the simulations was determined from Seahorse measurements. The oxygen concentration jumps between media channel and cell chamber are due to diffusion across the porous membrane. A dilute solution containing a small molecule entering the media channel at 20 μL/h diffuses across the porous membrane into the cell chamber and reaches a uniform concentration within the liver MPS within 300 seconds (right). Simulation assumes impermeable walls and no cell consumption. c) Preparation and microfabrication steps of the liver MPS. d) Photograph of a ready-to-use liver MPS. The catheter couplers link the tubes and microfluidic channels for perfusion. The short plug in the middle keeps the loaded cells in the chamber. e) Schematic of the liver MPS that shows the localization of the cell chamber; the red arrows indicate inlet and outlet of the media channels, whereas the black arrow marks inlet of the cell chamber. High cell viability in the tissue chamber after loading is demonstrated by fluorescence imaging of acetoxymethyl calcein (calcein-AM; live cells labeled green) and ethidium homodimer (dead cells labeled red). f) Immunofluorescent images of hiPSC-Heps seven days after loading into the liver MPS.

Oxygen consumption was an important factor in designing the liver MPS since hepatocytes have particularly high oxygen consumption rates (OCR). The performance of the liver MPS was simulated and optimized *in silico* using computational modeling (**Fig. 2b**; see Material and Methods for details). To maintain a high oxygen level in different flow conditions, the liver MPS was fabricated using a gas-permeable polydimethylsiloxane (PDMS). OCR in adult hepatocytes ranges from 0.3 to 0.9 nmol/s/10^6^ cells^51^, which is an order of magnitude higher than ~0.034 nmol/s/10^6^ in HepG2 cells^52^. The OCR in our hiPSC-Heps, measured using the Seahorse flux analyzer, was 0.1 nmol/s/10^6^ cells (**Supplementary Fig. 2**), which is consistent with previously reported results^53^. Using computational simulations, a media and cell chamber combination was designed that allowed for physiologically relevant oxygen levels in the cell chamber when considering the OCR of hiPSC-Heps and oxygen diffusion from the ambient through the PDMS walls. Using the OCR measured for our hiPSC-Heps, simulation showed a steady oxygen level of 0.16 mol/m^3^ across the device similar to the periportal region (pO_2_ 0.10-0.12 mol/m^3^ or 60-70 mmHg)^54^ after 24 hours (**Fig. 2b, left**). This tissue-level oxygen consumption is close to the physiological range and lower than hyperoxia (< 160 mmHg)^55^. Additional simulation assuming no oxygen diffusion through the PDMS showed oxygen depletion in the device after 300 seconds due to the cells’ high OCR (**Fig. 2b, center**). Shear stress, which is determined by the media flow rate used in the device, was lower than 5 dyn/cm^2^ previously found to diminish albumin and urea synthesis in rat pHeps^56, 57^. To give detail on the oxygen flux assumptions used in the simulation, additional COMSOL modeling information on the mesh and computational domains of liver MPS used are provided in (**Supplementary Fig. 3**). In our optimal design, it took an estimated 300 seconds using a 20 μL/h flow rate for a small molecule to reach a uniform concentration inside an empty device accounting for diffusion of the drug across the membrane into the cell chamber (**Fig. 2b, right**).

Fabrication of the liver MPS multilayer device was achieved with four parts: a glass slide, two layers of PDMS patterned with media and cell chambers, and a PET membrane. Media and cell chambers were permanently bonded into a device using oxygen plasma treatment and silanization for the PET membrane^58^ (**Fig. 2c**), creating the final liver MPS (**Fig. 2d**). The 3D liver microtissue in the MPS had a thickness of 80 μm covering the entire cell chamber of 5,560 μm (length) × 560 μm (width) in dimension. The liver MPS design permitted high cell viability after loading, indicated by live-dead staining of the cell chamber (**Fig. 2e**). Cellular function in the device was further confirmed by visualization of nuclear morphology, density and distribution of hiPSC-Heps, ALB and AFP expression, and formation of bile canaliculi (**Fig. 2f** and **Supplementary Fig. 4**). These assays showed that hiPSC-Heps were viable and maintained hepatocyte differentiation over 2 weeks of culture in the liver MPS.

### Comparison of synthetic function of hiPSC-Heps in liver MPS and conventional cell culture

Potential differences in differentiation and function between hiPSC-Heps in the 3D environment of the liver MPS and under 2D conditions in conventional cell culture dishes were assessed by analysis of albumin and urea secretion into the media. For this, hiPSC-Heps were released from the original differentiation cell cultures, split and either seeded into devices or replated on cell culture dishes. At the end of a 9-day time course, hiPSC-Heps secreted 41.41 +/− 3.53 μg/10^6^ cells/day albumin in the device, compared to 12.47 +/− 0.53 μg/10^6^ cells/day in conventional culture (Student’s t test, p=0.0036), a more than 3-fold increase (**Fig. 3a**). Similarly, hiPSC-Heps secreted 49.75 +/− 5.04 μg/10^6^ cells/day urea in the device, compared to 7.37 +/− 1.12 μg/10^6^ cells/day in conventional culture (Student’s t test, p=0.0009), a 7-fold increase (**Fig. 3b**). These results show that the liver MPS improves the synthetic function of hiPSC-Heps as compared to conventional cell culture.

**Fig 3.**
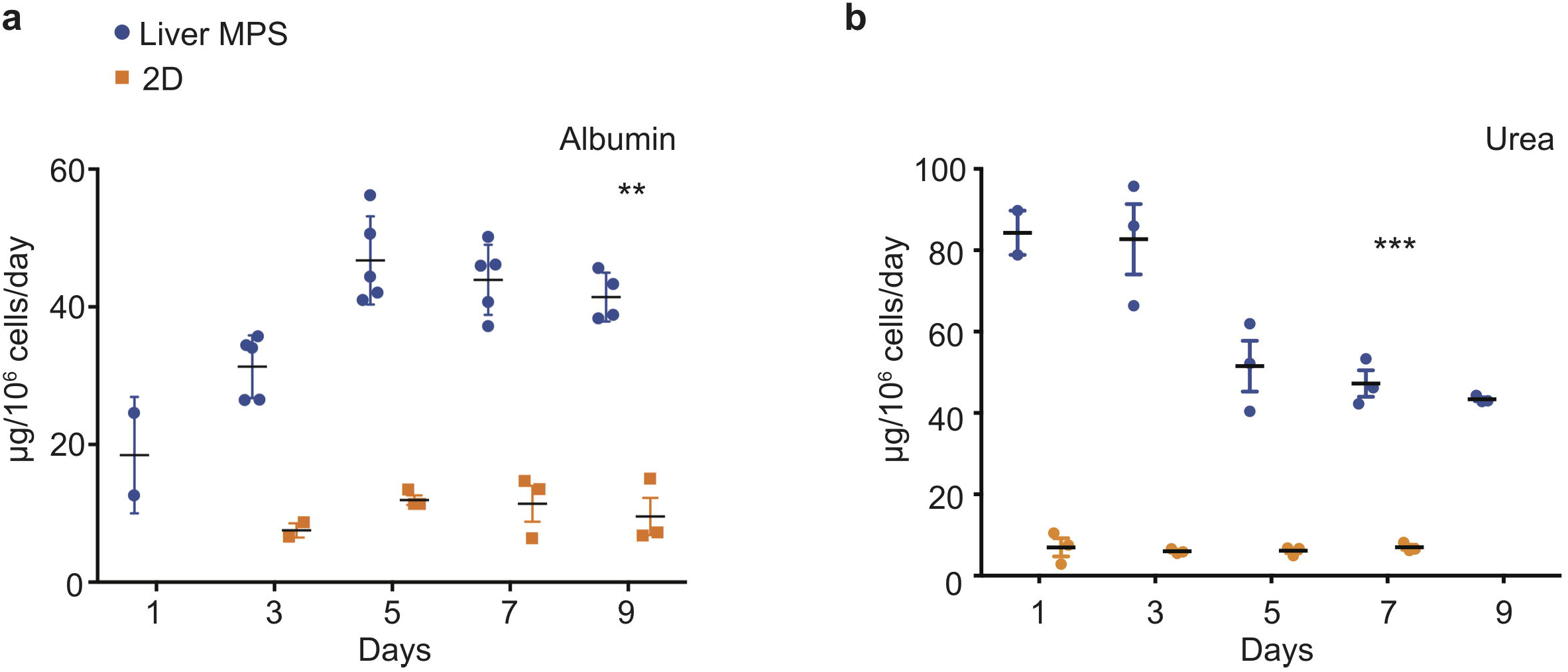
Synthetic function of hiPSC-Heps in conventional cell culture and liver MPS. a) Albumin secretion into the media measured on the indicated days after seeding hiPSC-Heps into the liver MPS or replating them on conventional cell culture dishes (2D). Data are presented as the mean +/− standard error of mean for five (liver MPS) and three (2D) independent experiments. Unpaired Student’s t-test with equal standard deviation, **p<0.05; ***p<0.01. b) Urea secretion into the media measured on the indicated days after seeding hiPSC-Heps into the liver MPS or replating them on conventional cell culture dishes. Data are presented as the mean +/− standard error of mean for three independent experiments. Unpaired t-test with equal standard deviation, ***p<0.01.

### Comparison of hiPSC-Hep-mediated cisapride metabolism in liver MPS and conventional cell culture

The arrhythmogenic drug cisapride is converted in pHeps into non-arrhythmogenic norcisapride^59^ by CYP3A4^60, 61^ (**Fig. 4a**). Therefore, the metabolism of cisapride can be inhibited or reduced by drugs and foods that act as CYP3A4 inhibitors such as ketoconazole or grapefruit (**Fig. 4a**). Cisapride metabolism in hiPSC-Heps was assessed in conventional cell culture using mass spectrometry to measure norcisapride formation within 30 minutes after addition of 1 or 10 μM of the parental drug to the cells. Intra- and extracellular amounts of norcisapride were measured and put in relation to the cisapride input concentration and the time the cells were exposed to the drug, allowing calculation of the metabolite formation clearance. Relative metabolite formation clearance for 1 μM of cisapride (100% +/− 10.81) was significantly reduced by addition of 10 μM of ketoconazole to 34.87% +/− 28.04) (**Fig. 4b**). Likewise, relative metabolite formation clearance for 10 μM of cisapride (100% +/− 6.08) was significantly reduced by addition of 10 μM of ketoconazole to 35.18% +/− 16.55) (**Fig. 4b**).

**Fig 4.**
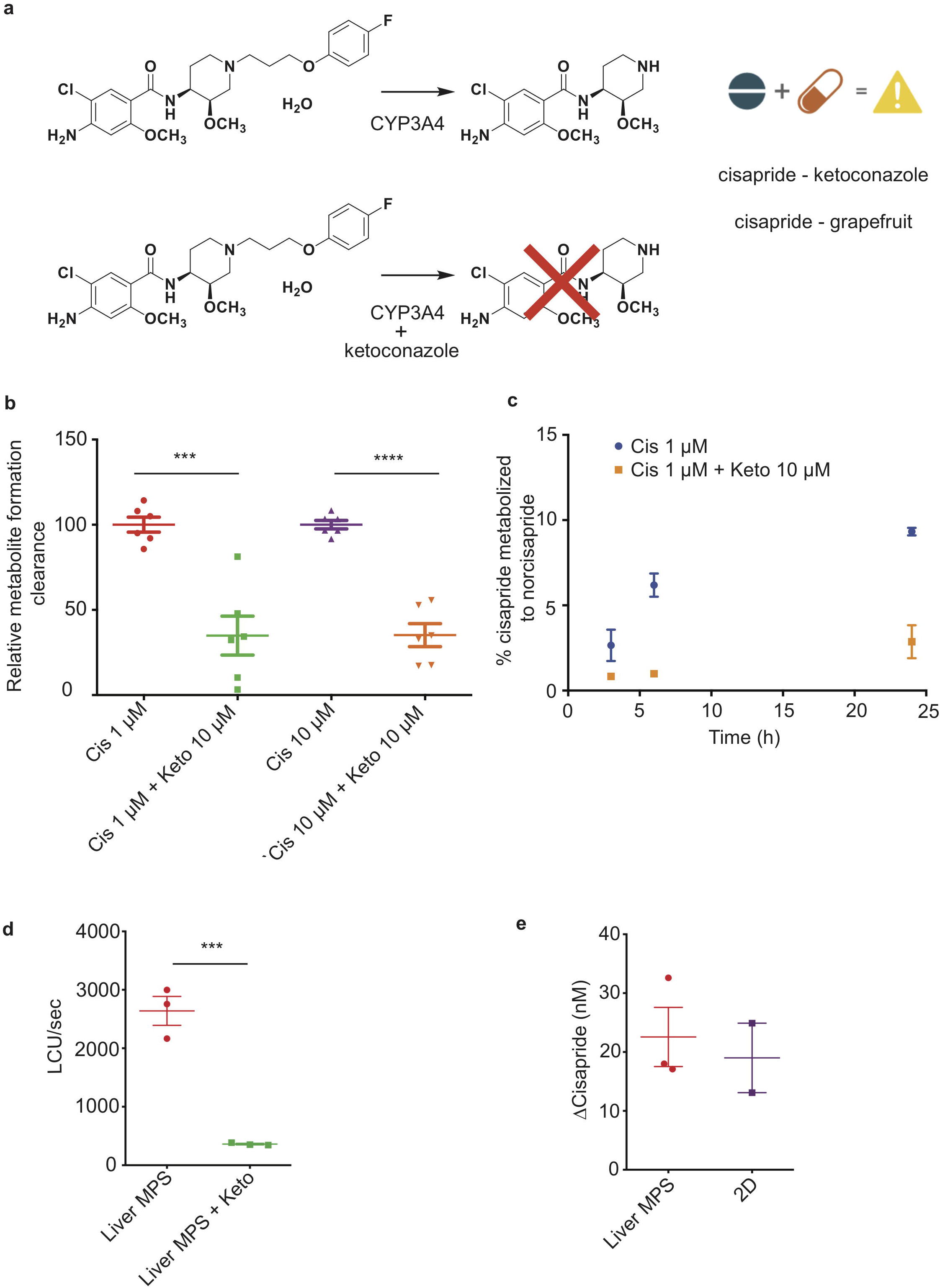
Cisapride metabolism in hiPSC-Heps in conventional cell culture and liver MPS. a) Metabolism of the arrhythmogenic drug cisapride into norcisapride by CYP3A4 in the liver (top). Drug-drug or drug-food interactions can inhibit this metabolic reaction. Ketoconazole (Keto) inhibits CYP3A4-driven metabolism of cisapride into norcisapride (bottom). b) Relative metabolite formation clearance for cisapride in hiPSC-Heps in conventional cell culture within 30 minutes. Data are presented as the mean +/− standard deviation from three independent experiments with each two biological replicates. Unpaired Student’s t-test, ***p<0.0003, ****p < 0.0001. Experiments were performed in cell culture dishes with low drug absorption. c) Percentage of cisapride metabolized to norcisapride with and without ketoconazole. Data are presented as the mean +/− standard error of mean for two biological replicates of one experiment. The experiment was performed in cell culture dishes with low drug absorption. d) CYP3A4 activity in the liver MPS measured with a luminescence assay with and without ketoconazole. Data are presented as the mean +/− standard error of mean for three independent experiments. e) Change in cisapride concentration in the liver MPS media channel and the supernatant of conventional cell cultures (2D). Data are presented as the mean +/− standard error of mean for three independent experiments.

To determine the ability of hiPSC-Heps to metabolize cisapride over an extended period of time, the percentage of cisapride converted into norcisapride was calculated 3, 6 and 24 hours after adding 1 μM of cisapride to the cells—without CYP3A4 inhibition by 10 μM of ketoconazole, 2.66 % (+/− 0.92), 6.19 % (+/− 0.69) and 9.33 % (+/− 0.23) of cisapride were metabolized, which dropped to 0.85% (+/− 0.02), 0.99% (+/− 0.06) and 2.87% (+/− 0.97) with ketoconazole, i.e., a reduction by 68, 85 and 70%, respectively (**Fig. 4c**).

Next, hiPSC-Hep-mediated cisapride metabolism was analyzed in the liver MPS. First, a non-destructive luminescence assay (Promega P450-Glo)—allowing serial analysis of samples taken from the media channel—was used to assess CYP3A4 activity based on cleavage-mediated activation of a luciferin substrate. As expected, additional infusion of 10 μM of ketoconazole into the device caused a 7-fold decrease in relative luminescence units (RLU) (2640 +/− 247 RLU vs. 361 +/−13 RLU; Student’s t test, p=0.0008) (**Fig. 4d**). Second, mass spectrometry was used to compare cisapride metabolism between hiPSC-Heps in the liver MPS or in conventional cell culture, which showed that cisapride concentrations were similarly reduced under both culture conditions 12 hours after exposure to 100 nM of the drug (**Fig. 4e**). These results establish that the small media volumes of the liver MPS are sufficient for luminescence assays and mass spectrometry, and that CYP3A4-mediated cisapride metabolism is maintained in hiPSC-Heps transferred into the device.

### Drug-drug interaction studies in integrated liver and cardiac MPSs

Before integration of liver MPS and cardiac MPS, cisapride-induced changes in beat rate and electrophysiology were analyzed in the cardiac MPS, which was created as previously described, including cardiomyocyte generation from the same hiPSC line used to generate hiPSC-Heps^4, 62^. In the clinical setting, the QT interval is used as a measure of cardiac electrophysiology. Since the QT interval depends largely on the heart rate, it is typically reported as QTc, a value normalized for 60 beats per minute (BPM) using Fridericia’s formula to introduce the heart rate correction^51, 63^. Optical measurement of action potential in the cardiac MPS^43^ was performed using the voltage-sensitive dye Berkeley Red Sensor of Transmembrane potential (BeRST-1; **Supplementary Fig. 5**)^64^. BeRST-1 is a far-red to nearinfrared dye that changes fluorescence intensity in response to membrane voltage changes through a photo-induced electron transfer mechanism. BeRST-1 dye was synthesized and its purity verified as previously described^64^. Using action potential duration at 80% of the repolarization (cAPD_80_) as a surrogate for QTc, an EC_50_=9.63 nM was observed in the cardiac MPS at cisapride concentrations ranging from 0 to 170 nM (after correction for absorption into the PDMS of the device) (**Fig. 5a**). Analysis of the spontaneous beat rate after direct application of escalating doses of cisapride (0 to 5 μM) to the cardiac MPS at 30-minute intervals showed a progressive BPM increase (**Fig. 5b**). Exposure of the cardiac MPS to 17 nM of cisapride caused a statistically significant prolongation of the QT interval (**Fig. 5c**).

**Fig 5.**
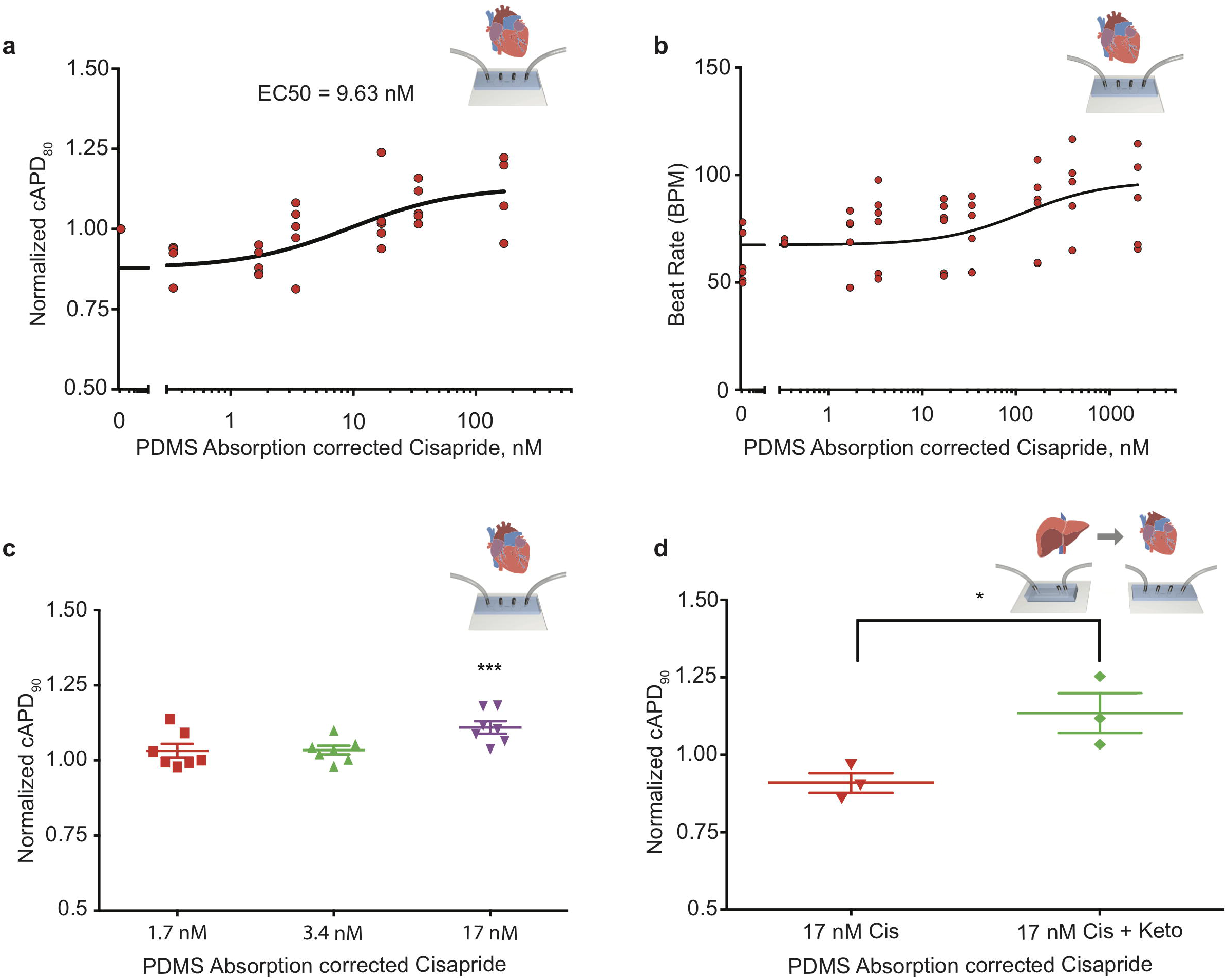
Cisapride effects on cardiac MPS and DDI in integrated liver and cardiac MPSs. a) Cisapride-induced changes in cAPD_80_ in the cardiac MPS. Non-linear regression fit log (inhibitor) vs. response (Hill Equation using three parameters) was used to obtain EC_50_ value. Cisapride values are corrected for PDMS drug absorption of 64%. Data are presented as the mean +/− standard error of mean for five independent experiments. b) Cisapride dose escalation study in the cardiac MPS. Spontaneous beat rate over dose range from 0 to 1 μM is shown. Cisapride values are corrected for PDMS drug absorption of 64%. c) Effect of increasing concentrations of cisapride on cAPD_90_ in the cardiac MPS. The cAPD_90_ values of the treatments were normalized the respective values for vehicle controls. Data are presented as the mean +/− standard error of mean for at least six independent experiments. Ordinary one-way ANOVA with Dunnett’s correction for multiple comparisons test, ***p < 0.0001. Cisapride values are corrected for PDMS drug absorption of 64%. d) Effect of cisapride metabolized in liver MPS on cAPD_90_ in cardiac MPS. cAPD_90_ values of the treatments were normalized to the 0 nM control values. Data are presented as the mean +/− standard error of mean for three experiments. Unpaired t-test with equal standard deviation was used, *p < 0.05. Cisapride values are corrected for PDMS drug absorption of 64%.

To facilitate study DDI of cisapride and ketoconazole and its effects on cardiac function, liver MPS and cardiac MPS were integrated. A prerequisite was the development of a common culture medium that supported the function of both hiPSC-Heps and hiPSC-derived cardiomyocytes in their respective devices. Baseline beat rate was functionally indistinguishable in RPMI plus B27-supplement or after 30-minute incubation in Williams E + Cocktail B media (**Supplementary Fig. 6**). For drug testing, cardiac MPSs were incubated with Williams E + Cocktail B media for 30 minutes and baseline cAPD_90_ values were recorded and normalized for each device. The cAPD_90_ in the cardiac MPSs was not altered by media from sequentially coupled liver MPSs perfused with 17 nM of cisapride for 8 hours. However, media from liver MPSs perfused with 17 nM cisapride together with 10 μM of ketoconazole for 8 hours significantly prolonged the cAPD_90_ in cardiac MPSs (**Fig. 5d**). These results reproduce in integrated hiPSC-derived liver and cardiac MPSs *in vitro* a DDI between cisapride and ketoconazole that was previously observed in patients *in vivo^65, 66^*.

## Discussion

This work focused on developing a functional liver MPS constructed from hiPSC-Heps, and using the system to study DDI-mediated inter-organ toxicity in a liver-cardiac MPS. There are two major reasons for creating better *in vitro* models of human hepatotoxicity and liver diseases that address the “fit for purpose” criteria used in the pharmaceutical industry. First, earlier and accurate safety profiling of a larger number of compounds, but still high throughput, can eliminate false positive attrition. Second, testing of more challenging hepatotoxicity mechanisms and complex disease models over longer time periods with lower throughput models representing physiologically relevant fluid flow and cell types^18, 67, 68, 69^. Focusing on the latter, we created a liver MPS employing hiPSC-Heps that provides a virtually unlimited source of patient-specific cells. Optimization of the differentiation protocol and in-depth characterization of the hiPSC-Heps showed functional drug uptake, metabolism and efflux systems, which make them suitable for use in drug screening and toxicity studies.

Microfabrication and microfluidic technologies provide a microenvironment that resembles *in vivo* conditions more closely than conventional cell culture^70, 71^. For example, in multi-well plates and culture flasks hepatocytes are cultured in monolayers with excessive medium volume-to-cell number ratios. More physiological medium-to-cell ratios can be achieved using the small system volumes of microfluidic devices^36^, which preserves the efficacy of low-concentration autocrine and paracrine signals, including secreted molecules, growth factors, matrices and hormones^72^. Along these lines, the microfabricated liver MPS described in this study can be used under a controlled range of flow rates, providing flexibility for optimizing cell viability and function. Using a PET membrane to mimic an endothelial cell barrier we protected the cells from flow-induced shear stress, which can compromise viability^56, 73, 74^. Moreover, under low shear stress conditions, perfusion/flow cultures provide better function of hepatocytes than static cultures, presumably due to greater nutrient exchange^75^. In accord, in our liver MPS high shear stress was only predicted in the inlet and outlet, which explains our finding of better function of hiPSC-Heps in the device than in conventional cell culture. Other factors promoting function of hiPSC-Heps in the device probably include 3D direct cell-cell contact and the low ratio of medium-to-cell volume. Whether the small subset of hiPSCs not committing to hepatocyte differentiation but giving rise to mesenchymal cell types instead (data not shown) further aided hiPSC-Hep function remains to be determined^76^.

The main goals for pharmaceutical companies in drug development are first safety, then efficacy. For go/no-go decisions to move a drug down the pipeline a safety margin is employed, which is calculated from the drug’s estimated therapeutic plasma concentration (ETPC) and the *in vitro* EC_50_ or IC_50_ (e.g., EC_50_/ETPC). Using our cardiac MPS we evaluated the *in vitro* EC_50_ of cisapride to be 9.63 nM when cAPD_80_ was used a metric. This EC_50_ value is relatively close to previous estimates of 32 nM^77^ in hiPSC-derived cardiomyocytes and 21.2 nM^78^ or 20 nM^79^ in human HEK293 cells (**Supplementary Table 1**). Based on the ETPC of cisapride being 2.6-4.9 nM^80^, we calculated a margin of safety of approximately 3.7-fold, which is close to the clinically determined safety margin of 1.1 to 6.5-fold^80, 81, 82^. Redfern *et al*.^80^ have suggested that a margin of 30-fold provides a degree of safety necessary for drugs that can cause cardiac arrhythmias. The data presented suggests the use of our liver-cardiac MPS would have correctly predicted cisapride’s negative effects.

Our liver-cardiac MPS successfully replicated a highly publicized example of DDI severely impacting human health despite successful clinical trials. Cisapride, a prokinetic used to treat gastroesophageal reflux^46^, causes prolongation of the QT interval, ventricular arrhythmias and torsade de pointes^47^ if metabolism of the drug is impaired. In humans, cisapride is extensively metabolized to inactive norcisapride through N-dealkylation (41%-45% of the administered dose) and to several minor metabolites^60^. CYP3A4 is involved in the metabolism of most drugs in the human liver, including cisapride N-dealkylation^59, 83, 84^. DDI with ketoconazole or drug-food interactions (e.g., grapefruit), leading to inhibition of CYP3A4, caused 23 deathsand 341 incidences of torsade de pointes, and ultimately removal of cisapride from the market^48, 49^.

We used this example to demonstrate the robustness of liver-cardiac MPS interaction studies to predict DDI. First, we validated the cisapride metabolism of hiPSC-Heps using LC-MS/MS and a luminescence assay and confirmed that CYP3A4 metabolic function was inhibited by the addition of ketoconazole to the media using the norcisapride metabolite as a reference value (**Supplementary Fig. 7**). For this analysis, we corrected the observed QT interval using Fridericia’s formula for rate correction of QT interval^51, 63^. Then we confirmed prolongation of the normalized corrected (cAPD_90_) interval in the cardiac MPS when exposed to media retrieved from the liver MPS in the presence of cisapride and ketoconazole. Next, we analyzed cisapride’s dose-escalation effects on the cardiac MPS and determined cisapride concentrations that affected the cAPD_90_. The effect was eliminated in the sequentially functional coupled liver-cardiac MPS where cisapride was first passed through the liver MPS prior to reaching cardiac MPS.

The ability to predict DDI validates the function of our liver-cardiac MPS, and illustrates its ability to predict unforeseen medical complications. The integrated system exhibits a level of biological complexity that approaches *in vivo* conditions, thereby providing a powerful platform for simultaneously screening drug efficacy and toxicity. Although differentiation and function of hiPSC derivatives generated with current protocols are not equal to their primary counterparts, our study shows that in-depth functional characterization allows using these cells to build genetically defined faithful multi-organ disease models.

## Materials and Methods

### Cell Sources

The hiPSC line WTC was generated by reprogramming of skin fibroblasts derived from a healthy 30-year-old Japanese adult male with no known family history of heart disease, generously provided by Prof. Bruce Conklin (Gladstone Institutes) and available from the Coriell Repository (#GM25256). WTC was used for generation of hiPSC-Heps. hiPSC-derived cardiomyocytes were generated from WTC edited to harbor a single-copy of CAG-driven GCaMP6f knocked into the first exon of the AAVS1 “safe harbor” locus^85^. For experiments with pHeps, commercially available cryopreserved cells from ThermoFisher Scientific (lot#Hu8138) were used.

### Differentiation of hiPSC-Heps

Differentiation of hiPSCs (day 0) into hiPSC-Heps (day 23 – 25) was performed according to an optimized protocol. Cells were cultured on Cultrex Basement Membrane Extract (BME); (Trevigen) diluted 40 times in Knockout DMEM (ThermoFisher Scientific) for the entire hepatocyte differentiation process. At day 0, hiPSCs were split and kept in mTESR (Stem Cell Technologies) supplemented with 10 μM Rock inhibitor (Tocris Bioscience) to form a confluent layer of hiPSCs on day 1 when the hepatocyte differentiation was initiated. From day 1 to 7, RPMI 1640 medium (ThermoFisher Scientific) was used and supplemented with standard antibiotics and antimycotics and recombinant human Activin A (PeproTech EC) at 100 ng/mL. Between day 2 and 7, Gem21 NeuroPlex without Insulin (Gemini BioProducts), non-essential amino acids (ThermoFisher Scientific) and sodium butyrate (Sigma Aldrich) at 500 μM was added. KnockOut Serum Replacement (ThermoFisher Scientific) was added to the differentiation medium from day 1 to 3 in decreasing amounts of 2, 1 and 0.2%, respectively. The small molecules CHIR 99021 (Tocris Bioscience) at 3 μM and PI-103 (Selleckchem) at 50 nM were added at day 1 and from day 1 to 3, respectively. From day 8, cells were cultured in IMDM (ThermoFischer Scientific) supplemented with standard antibiotics and antimycotics, Gem21 NeuroPlex without Insulin (Gemini BioProducts), 100 nM insulin, 100 nM dexamethasone and 400 μM monothioglycerol (all Sigma Aldrich), non-essential amino acids and human FGF2 at 10 ng/mL and human BMP4 at 20 ng/mL (both Peprotech) until day 18. At day 13 HGF (Peprotech) at 20 ng/mL was added until the end of the hepatocyte differentiation. From day 18 onwards, cells were cultured in Hepatocyte Basal Medium (Lonza CC-3199) supplemented with all components of Hepatocyte Culture Medium SingleQuots except EGF (Lonza CC-4182 containing gentamicin-amphotericin, transferrin, insulin, hydrocortisone and fatty acid-free bovine serum albumin). This medium is referred to as Hepatocyte Culture Medium (HCM).

### Immunofluorescence in conventional cell culture and liver MPS

In conventional cell culture, hiPSCs and hiPSC-Heps were fixed for 20 minutes at room temperature (RT) in 4% paraformaldehyde (PFA) and washed 3 times with phosphate-buffered saline (PBS). Cells were blocked for 1 hour with 3% BSA and permeabilized with 0.3% Triton X-100 (Sigma). Cells were labeled with rabbit anti-OCT3/4 antibody (sc-9081; SantaCruz Biotechnology) diluted 1:500, goat anti-HNF4 antibody (sc-6557; SantaCruz Biotechnology) diluted 1:500, goat anti-human albumin antibody (A80-129A; Bethyl Laboratories) diluted 1:500, mouse anti-alpha-fetoprotein (MA1-19342; Invitrogen) diluted 1:500 and DAPI (D1306; Invitrogen) for nuclear staining. Secondary antibodies were Alexa Fluor 488/555 donkey-anti rabbit or anti-goat or anti-mouse IgG diluted 1:1000 (Invitrogen). Cells were imaged using a Nikon Eclipse TE300 with a Lumencore Spectra X light engine. Liver MPS were washed with PBS and fixed at RT for 30 minutes with 4% PFA using a 3D collagen gel staining protocol and washed 3 times with PBS. After permeabilization in 1% Triton X-100 (Sigma) for 30 minutes, cells were incubated for ~24 hour at 4°C with the same primary antibodies as above but at different concentrations: Both goat antihuman albumin and mouse anti-alpha-fetoprotein were diluted 1:100. Secondary antibodies with fluorescent probes Alexa 488 (A-11055; Invitrogen) and R-PE 565 (P21129; Invitrogen) were incubated at 4°C for 6 hours and washed 3 times for 5 minutes with cold PBS. Liver MPS were imaged by confocal microscopy. Confocal images were collected using a Carl Zeiss LSM 710 confocal microscope equipped with a plan-apochromat 10Å~/0.45 objective.

### Flow cytometry

At day 23 of differentiation, hiPSC-Heps were digested with 1 mg/mL collagenase for 20 minutes to singularize cells. Cells were then fixed with 4% PFA for 20 minutes at room temperature and labeled with the following antibodies: Anti-human albumin (A80-129A; Bethyl Laboratories) detected with a secondary antibody conjugated with Alexa-488 (Life Technologies) and anti-ASGR1 (8D7 clone conjugated with PE; BD Biosciences). Albumin and ASGR1 positive cell populations were determined by flow cytometry analysis with an Attune NxT Acoustic Focusing Cytometer (Thermo Fisher Scientific). FlowJo software was used for the analysis of the data.

### Drug transport study

Cells were washed with carbogenated William’s E (Gibco; A1217601) (adjusted pH 7.4 at 37°C) for 15 minutes at 37°C before transport or metabolism studies. The drug uptake transport study was initiated by adding each transporter substrate with or without its inhibitor in William’s E medium, and incubated at 37°C for 2 minutes in a shaker-incubator. The reaction was terminated by removing the dose solution followed by washing the cells 3 times with ice-cold PBS. The following substrates and inhibitors were used: 10 μM acyclovir (including 0.1 μCi [^3^H]-acyclovir, OAT substrate) and 1 mM probenecid (OAT inhibitor) for OAT transport study, 1 mM metformin (including 1 μCi [^14^C]-metformin, OCT substrate) and 1 μM decynium-22 (OCT inhibitor) for OCT transport study and 10 μM rosuvastatin (including 0.25 μCi [^3^H]-rosuvastatin, OATP substrate) and 50 μM rifamycin-SV (OATP inhibitor) for OATP transport study. Cells were harvested for further drug analysis. The drug efflux transport study was initiated by adding each transport substrate with or without its inhibitor in William’s E medium and incubated at 37°C for 15 minutes in a shaker-incubator. The reaction was terminated by removing the dose solution followed by washing cells 3 times with ice-cold PBS. The following substrates and inhibitors were used: 10 μM digoxin (including 0.25 μCi [^3^H]-digoxin, P-gp substrate) and 10 μM GG918 (P-gp inhibitor) for P-gp transport study and 10 μM of mitoxantrone (including 1 μCi [^3^H]-mitoxantrone, BCRP substrate) and 5 μM GG918 (BCRP inhibitor) for BCRP transport study. Cells were harvested for further drug analysis (see ‘Drug metabolism measurement’ section).

### CYP luciferase activity assay

CYP activity was assessed using the luciferin-based P450-Glo assays (Promega). Cells were washed with basal HCM for 15 minutes prior to incubation with CYP substrates at the following concentrations: 100 μM for 1A2, 100 μM for 2C9, 10 μM for 2C19, 30 μM for 2D6, 3 μM for 3A4 and 150 μM for 3A7. Cells were incubated with substrates for 3.5 hours except for the 3A4 assay that was incubated for 70 minutes. At the end of the incubation period, 50 μL supernatant was transferred into a 96-well plate in technical triplicates, and 50 μL of CYP-specific detection reagent was added to each well. The plate was covered in foil and mixed gently for 20 minutes, and absorbance was measured at 450 nm on a plate reader (SpectraMax i3, Molecular Devices) Samples included a negative control consisting of HCM only (no cells) that was used for background correction.

### Drug metabolism enzymatic assays

Each enzyme substrate was dissolved in William’s E medium and incubated at 37°C in a shaker-incubator. For metabolism studies of UGTs and SULTs, cells were incubated with 10 μM 1-naphthol or 10 μM nitrophenol (Sigma-Aldrich), respectively, for 15 minutes at 37°C. Then the dosing solution was collected, the cells were washed 3 times with ice-cold PBS and harvested for further drug analysis (see ‘Drug metabolism measurement’ section).

### qPCR

RNA was extracted from hiPSC-Heps and pHeps by the Trizol/chloroform method. Each sample was suspended in 500 μl of Trizol to which 100 μl (1V:5V) of chloroform were added. After 15 seconds of vortexing and 5 minutes of incubation at RT, samples were centrifuged for 5 minutes at 12,000 rpm. The upper phase containing RNA was transferred into a new Eppendorf tube. Isopropanol (1V: 1V) was added to each tube and slowly inverted five times. Samples were centrifuged for 10 minutes at 12,000 rpm at 4°C. The RNA pellet was kept on ice until the end of the procedure. The pellet was washed twice in 750 μl of 70 % ethanol and air-dried on ice. The pellet was resuspended in an appropriate volume of nuclease-free water and the RNA concentration was determined on a Nanodrop (ThermoFisher Scientific). Reverse transcription was performed with 1 μg of RNA template using cDNA Supermix (Quanta Biosciences) and nuclease-free water in a final volume of 20 μl. qPCR was performed on 100 ng of cDNA using SYBR Green Supermix (Affymetrix) and specific primers on an Applied Biosciences ViiA7 Real-Time PCR System (Invitrogen).

### Computational modeling

Laminar fluid flow and dilute species transport in the liver MPS was simulated using COMSOL Multiphysics 5.3a (COMSOL AB). Laminar fluid flow was governed by the Navier-Stokes equations and dilute species (oxygen (O_2_) and a generic small molecule) transport by the advection-diffusion equation, both solved in COMSOL subject to the following boundary conditions: No-slip on all media channel walls and membrane surfaces; laminar inflow rate specified at the media channel inlet; laminar outflow conditions at the outlet; dilute species concentration specified at inlet. O_2_ diffusion through the 3.5-mm thick PDMS slab was also simulated in COMSOL, subjected to ambient O_2_ concentration at the top of the slab and O_2_ partial pressure continuity at the media channel and cell chamber walls. In our simulations, fluid (media with supplements) with a density of 1000 kg/m^3^ and a dynamic viscosity of 0.78 mPa·s at 37°C^86, 87^ was used. Flow across the 15-μm thick porous membrane was neglected since low membrane porosity (5.6%), small membrane pore size (mean pore diameter 3 μm) and cultured cells on the cell chamber side of the membrane resulted in flow resistances through the membrane that were orders of magnitude higher than through the media channel (height 100 μm)^58^. Moreover, the cultured cells within the cell chamber increased flow resistance there. Thus, it was assumed that there was no media flow into or within the cell chamber so that dilute species transport within the cell chamber was by diffusion only^58^. Small flows within cell culture chambers in membrane bilayer microfluidic devices have been considered elsewhere^88, 89^.

Three sets of COMSOL simulations were conducted. In the first set, the transport of a generic small molecule within culture media in the liver MPS was modeled with an inlet flow rate of 20 μl/h, an inlet concentration of 1 mol/m^3^, zero initial concentration in the device, a diffusion coefficient of 1.0 × 10^−9^ m^2^/s in culture media and no cell uptake and no flux through walls. The second and third sets of simulations involved O_2_ transport, diffusion and uptake. The media channel inlet and initial media channel and cell chamber O_2_ concentrations were 0.173 mol/m^3^, which is the saturation level in culture media at 37°C in equilibrium with ambient incubator air at 37°C, 1 atm pressure, with 18.7% O_2_ (21% O_2_ in dry air reduced by 5% CO_2_ and 6% water vapor). On the PDMS surfaces exposed to ambient incubator air, O_2_ concentration in the PDMS was 1.11 mM, the saturation level. Diffusion coefficients of O_2_ in PDMS and culture media were 3.25×10^−9^ m^2^/s and 3.0×10^−9^ m^2^/s, respectively^87, 90, 91^. Cell oxygen consumption was modeled by Michaelis-Menten kinetics with rate = −VO_2max_ • ρ_cell_ • c/(K_m_ • S_cell_+c), where c was the local oxygen concentration, K_m_ = 5.6 mmHg is the Michaelis–Menten constant^54, 92^, and S_cell_ = 1.049 mM/(atm O_2_) was the solubility of oxygen in cells^92, 93^. VO_2max_ = 1.04×10^−16^ mol/s/cell was measured for hiPSC-Heps using the Seahorse XF24 cellular respirometer (Agilent) (see, Oxygen Consumption Rate section for details). A cell density in the cell chamber of ρ_cell_ = 6.44×10^13^ cells/m^3^ based on ~ 10,000 (9812 +/838) cells was assumed per liver chip.

### Liver MPS fabrication and design dimensions

The liver MPS was designed to mimic a ~100 parallel circuit of liver sinusoids (~6-15 μm in width)^94, 95^. The design consisted of a rectangular chamber divided by a PET membrane to mimic the fenestrations present in the sinusoidal endothelial cells in the human liver^96^. The liver MPS was composed of two chambers with the following dimensions: 5,560 μm (length) × 560 μm (width) × 100 μm (height) for the cell chamber side, and in the media channel the volume was 0.32 μL. The total volume of the chip was empirically measured to account for the cell inlet and media outlet having a total volume of 2 μL. Computational modeling (see previous section) was used to verify the chip dimensions and fluidic design were adequate for transport of O_2_ and drugs.

The dual chamber microfluidic device was fabricated using replica molding from photolithographically defined SU-8 masters on silicon wafers. Briefly, 30 g (top layer) and 6 g (bottom layer) of PDMS was poured into two silicon wafer masters and baked at 80°C for 8 hours, after which O_2_ plasma was used to bond PDMS devices to the PET membrane with 3 μm pore size (Sabeu GmbH) and the cover glass. To adhere the PET membranes, they were treated with bis[3-(trimethoxysilyl)propyl]amine silane and then rendered hydrophilic (protocol modified from Sip *et al*.^58, 91^). This treatment also allows for PET to remain bonded even in the presence of salt-containing media. The PET membranes were prepared by first being cut into slightly larger rectangle shapes than those of the individual cell or media layer patterns and then cleaned in isopropyl alcohol for 10 minutes. The membranes were then suspended in a plasma chamber that was filled with 20% O_2_ and exposed to plasma at 60W for 60 seconds. The power source used was a PE-1000 AC Plasma power source. The membranes were then incubated in a solution of 97% isopropyl alcohol, 2% bis(3-(trimethoxysilil)propyl)amine and 1% H_2_O at 80°C for 20 minutes. Following incubation, the membranes were rinsed with isopropyl alcohol and dried at 80°C for 30 minutes. Following drying (*curing*), the membranes were individually placed in 2 mL of 70% ethanol in type I ASTM laboratory reagent grade H_2_O for 30 minutes to render the membrane hydrophilic.

### Liver MPS loading with hiPSC-Heps

Prior to cell loading, the MPS was sterilized by UV for 30 minutes, coated with 100 μg/mL human fibronectin (Corning, USA) and 100 μg/mL rat tail collagen, type I, alpha 2 (Corning, USA) in PBS, and the interior of the device was dried under vacuum for 30 seconds. hiPSC-Heps were trypsinized for 8-10 minutes and quenched using 80% DMEM/F12 (Thermofisher, Carlsbad, CA) with 20% FBS, pelleted at 50 g for 5 minutes, and then resuspended at 11 × 10^6^ hepatocytes/mL in HCM supplemented with 10% fetal bovine serum (Cellgro) and 10 μM Rock inhibitor. hiPSC-Heps were then pipetted into the cell inlet port of the device and centrifuged in a bucket at 300 × g for 3 minutes for optimal cell loading. Seeded devices were incubated overnight at 37°C to enhance the formation of a 3D organization. Plating media was removed from the device and HCM was added and incubated for 24 hours at 37°C to allow stabilization of the model before initiating a 10 μL/h flow perfusion using a syringe pump (New Era NE-1800).

To quantify the number of total cells in the liver MPS, confocal images of cell nuclei stained with DAPI in the cell chamber were imaged by z-stack confocal microscopy (Zeiss LSM 710). Then, the Cell Counter plugin of Fiji ImageJ was used to manually count the cells as the automated counter was not able to distinguish cells accurately in 3D. This process was repeated in triplicates and a best-estimate of the average cell number (9812 +/− 838 cells) in the liver MPS was used to normalize albumin and urea secretion per cell.

### Live/dead assays

To assess viability of hiPSC-Heps in the liver MPS, the LIVE/DEAD Cell Imaging Kit from Molecular Probes (R37601) was used. Viable cells were stained with green-fluorescent calcein AM (488 nm) and dead cells with red ethidium homodimer-1 (570 nm). The liver MPS was rinsed with PBS and 60 μL of dye were added via the media channel and incubated for 20 minutes. Images were collected using an inverted Nikon TE300 microscope with a Lumencore Spectra X light engine. Images were processed with Fiji ImageJ. Counts of live or dead cells were obtained using a segmentation analysis coupled to a constant area exclusion filter. For long-term culture, the liver MPS was incubated at 37°C and fed by a New Era NE-1800 syringe pump with continuous media flow (10 μL/h). For visual characterization, cells were imaged using a brightfield phase contrast daily without detachment from the syringe pump using a Nikon Eclipse TE300 microscope.

### Efflux media collection and biochemical measurements in liver MPS

Perfusion of the liver MPS was initiated (day 0) 2 days after loading to allow recovery and spreading of the hiPSC-Heps. Media was collected every two days and analyzed for albumin and urea secretion. Albumin was measured using an enzyme-linked immunosorbent assay (Bethyl Laboratories). Urea nitrogen was measured using a colorimetric assay (Urea Nitrogen Test, Stanbio Laboratory) modified to be performed in a 384-well microtiter plate, including increase of the incubation time of reactants from 60 to 90 minutes at 60°C prior reading. All biochemical assays were performed in 10 μL of media and measured on a SpectraMax i3 plate reader (Molecular Devices). All sample results were calculated by interpolation of sample raw values from standard curves performed in parallel. The same assays were repeated for conventional cell cultures of hiPSC-Heps.

### Drug metabolism of hiPSC-Heps in conventional cell culture and liver MPS

For drug metabolism study in conventional cell culture, hiPSC-Heps were incubated with 1 or 10 μM of cisapride (Sigma) in Williams E + Cocktail B (see below) with or without 10 μM ketoconazole (Sigma-Aldrich) for the indicated time-points (from 30 minutes to 24 hours) at 37°C in an incubator. Then the dosing solution was collected, the cells were washed 3 times with ice-cold PBS and harvested for further drug analysis by mass spectrometry (see ‘Drug metabolism measurement’ section). To determine CYP3A4 activity and its inhibition by ketoconazole in the liver MPS after day 25, a luminescence assay (P450Glo, Promega) was used. Briefly, the liver MPS was incubated with Luciferin-PFBE for 24 hours, after which the supernatant was transferred to a 96-well plate and luciferin detection reagent was added to initiate a luminescent reaction. The plate was then incubated for 20 minutes at RT to complete the reaction. Luminescence levels were measured on a SpectraMax i3 plate reader. CYP3A4-mediated metabolism of cisapride was performed in Williams E + Cocktail B. Media was supplemented with 100 nM and 1,000 nM cisapride and perfused through the liver MPS on differentiation day 26. These concentrations were subsequently corrected for absorption into PDMS (see, below). Efflux media was collected over a 24-hour period and stored at −80°C for further analysis by mass spectrometry (see ‘Drug metabolism measurement’ section).

### Drug metabolism measurements

The cisapride drug and norcisapride metabolite were extracted from efflux media and analyzed using a Shimadzu UFLC system (Carlsbad) coupled with a Sciex API5000 triple quadrupole mass-spectrometer (Foster City) using positive ionization mode. Cisapride and norcisapride were further separated by a gradient mode of 10 mM ammonium acetate-acetonitrile as mobile phase using Acclaim Trinity P1 reverse phase column (2.1 × 50mm, 3 μm; Thermo Fisher Scientific). Mass ion transitions (Q1/Q3) of cisapride and norcisapride were *m/z* 468.3/186.0 and *m/z* 314.2/184.0, respectively. 1-naphthol, 1-naphthol-s-glucuronide, *p*-nitrophenol and *p*-nitrophenol sulfate were separated by a gradient elution with 0.1% formic acid in H_2_O and 0.1% formic acid in acetonitrile as mobile phases using a XTerra MS C18 reverse phase column (4.6 × 50 mm, 3 μm; Waters). Mass ion transitions of 1-naphthol, 1-naphthol-s-glucuronide, *p*-nitrophenol and *p*-nitrophenol sulfate were *m/z* 143.3/115.3, *m/z* 318.9/143.3, *m/z* 138.2/168.1 and *m/z* 217.9/138.2, respectively. Standard curve range of each drug and metabolite was 0.5-2,000 nM. Precision (defined by the coefficient of variation) and accuracy (defined by relative error) of LC-MS/MS analyses were both <15% for all drugs and metabolites.

### Cisapride EC_50_ studies

Culture medium for the cardiac MPS was RPMI 1640 medium (11875-093; Gibco) supplemented with B-27 (17504-044; Gibco). The cardiac MPS was generated as described in a previous study^4^. For all pharmacology, the cardiac MPS was first equilibrated to phenol red free Williams E media + Cocktail B (see below) containing vehicle control (DMSO or water, to a final concentration of 0.1% v/v). On the day of the experiment, freshly measured cisapride drug was dissolved into DMSO.

After the initial baseline recording in vehicle control condition, media were exchanged for the lowest cisapride drug dose, and the cardiac MPS was incubated for 30 minutes at 37°C. Media changes during the drug testing were either performed manually^62^ or using a Fluigent pump system. A volume flow of 30 μL/min was applied for 5 minutes to change to a new drug dose followed by 15 μL/min continuous perfusion. Spontaneous beating activity was recorded for brightfield, GCaMP and BeRST imaging analysis as described below. For action potential recording, cardiac MPS were first labeled overnight with 2.5 μM BeRST-1. For the cardiac MPS metrics analysis, cAPD_80_ and cAPD_90_ were used which is the beat-rate corrected action potential duration at 80 and 90% repolarization (see example in **Supplementary Fig. 5b**).

### Cisapride drug testing in cardiac and liver MPS

One initial approach to coupling organs is direct coupling of the media outlet from one organ into the inflow of the next. This approach requires more refinement due to the complexity needed for a common medium in aspects such as reducing metabolic wastes to the next organ, organ-specific growth factors and flow rates, and adequate levels of O_2_ throughout the organs in the system. Functional coupling of the liver and cardiac MPS started with testing of the basal pHep maintenance William’s E medium in the cardiac MPS to determine effects on electrophysiology and toxicity of hiPSC-derived cardiomyocytes prior to DDI studies. William’s E medium was supplemented with Gibco Cell Maintenance Supplement Pack B (Cocktail B; Thermofisher Scientific), which contains bovine serum albumin (5.35 μg/mL) as a carrier protein and other supplements such as ITS that support pHep culture. Dexamethasone was not included to exclude likely confounding effects on hiPSC-derived cardiomyocytes.

Before integration, cardiac and liver MPSs were equilibrated in William’s E medium supplemented with Cocktail B containing vehicle control DMSO at a final concentration of 0.1% v/v. Cisapride was infused into the devices as a 0.1% DMSO solution in William’s E medium + Cocktail B. On the day experiments were performed, freshly prepared concentrated stock (10 mM) of cisapride (in DMSO) was dissolved into William’s E + Cocktail B and then diluted to working concentrations (nM range). After recording activity in zero-dose vehicle condition, media were exchanged for the lowest drug dose, and MPSs were incubated for 30 minutes at 37°C for the dose escalation studies. Spontaneous beating activity of hiPSC-derived cardiomyocytes was recorded as described above. This was repeated for each dose of drug. For the media transfer experiments, 100 μL of cisapride were incubated in the liver MPS for 8 hours. After this incubation time, the media (referred to as cisapride conditioned media) was stored at −20°C for future usage in the cardiac MPS.

### Cardiac MPS imaging and analysis

All images were collected using a Nikon Eclipse TE 300 microscope. Brightfield videos were analyzed for beating physiology using an updated version of our open source motion tracking software^85^. The software can be downloaded at https://sites.wustl.edu/huebschlab/resources/. Microscopy files were directly read into the MATLAB-based graphical user interface (GUI), and the contractile motion was analyzed via an exhaustive-search block-matching optical flow algorithm that compared the position of 8×8 pixel macroblocks at frame *i*to their position at frame *i*+5 (corresponding to the motion in 50 msec). Motion vectors were used to calculate beat-rate.

BeRST-1 data were quantified using in-house MATLAB code that was developed based on previous work by Laughner and Bogess et al^98, 99^. BeRST-1 videos were analyzed for beat rate and 80% and 90% action potential duration (APD_80_, APD_90_), using custom MATLAB and Python scripts (**Supplementary Fig. 5**).

### Characterization of drug absorption to PDMS

While PDMS is widely used to build microfluidic devices, due to its biocompatibility and convenience for microfabrication, the material has the disadvantage that it can absorb small molecules with low LogP values^100, 101^. Thus, absorption of cisapride to PDMS was characterized to correct the dosages used in the experiments. Dose escalation studies were first repeated in the liver MPS, without cells, to model the drug analysis. Using LC-MS/MS, ion abundances of the drug in stock samples and in efflux media were measured to calculate the percent remaining. One device was used per escalation study of 10, 50, 100, 500 and 1,000 nM, and the average percent remaining were 35.52, 30.54, 30.95, 34.55 and 39.79 nM, respectively (**Supplementary Fig. 8**). For the drug response studies, the cisapride concentration was corrected for PDMS drug absorption (60-70%)^100, 102^.

Assembled liver and cardiac MPS chips were placed on a temperature-controlled glass base (Thermo Plate; Tokai Hit) to maintain the temperature of the chips at approximately 37°C. The drug cisapride was prepared to the concentrations of 5, 10, 50 and 100 nM in William’s E medium supplemented with Cocktail B. The MPSs were loaded with 100 *μ*L of drug solution via a pipette tip at the media inlet port and an empty pipette tip placed at the media outlet. Flow at the media channel was induced due to the difference in hydrostatic pressure head at the inlet and outlet. For a given concentration of drug, the system was given 30 minutes after loading before the pipette tip at the inlet was taken out and the solution in the device and at the outlet tip was collected. Experiments were started with the lowest concentration (i.e., 5 nM) and dose concentration was increased for subsequent experiments (i.e., dose escalation). Technical replicates for each drug concentration are indicative of solutions collected from different devices. Drug solutions collected from devices were diluted by half using a 10 nM solution of propranolol used as an internal standard for LC-MS/MS. Standards of cisapride drug with propranolol as internal control were also prepared. Samples were analyzed using a LC system (1200 series, Agilent) connected in line with an LTQ-Orbitrap-XL MS equipped with an electrospray ionization source and operated in the positive ion mode (ThermoFisher Scientific). The LC was equipped with a reversed-phase analytical column (length: 150 mm, inner diameter: 1.0 mm, particle size: 5 μm; Viva C18; Restek). Acetonitrile, formic acid (Optima LC-MS grade, 99.5+%; Fisher), and water purified to a resistivity of 18.2 MΩ·cm (at 25°C) using a Milli-Q Gradient ultrapure water purification system (Millipore) were used to prepare mobile phase solvents. Data acquisition and processing were performed using Xcalibur software (Version 2.0.7, ThermoFisher Scientific). Processed data was used to quantify the amount of drug lost due to absorption in the PDMS-based MPSs via comparison with the drug standards.

### Statistical Analysis

Differences in drug and control population measurements reported as significant have a p < 0.05, as determined by application of a t-test with the assumption of equal variance. Direct comparisons were made by non-paired Students t-test and for multiple comparisons ANOVA analysis was used. All curve fitting was done using GraphPad Prism 6.

## Authors’ contributions

All authors participated in the study design, analysis of the data, interpretation of the results and review of the manuscript; FTLM, AL, LD conducted the main experiments; AL and HO analyzed drugs and metabolites; BS helped with cell culture; VC helped with cardiac MPS drug testing; IG and CL helped with computer modeling, drug absorption studies and liver MPS analysis; SB and EM provide BeRST; and FTLM, AL, HW, KEH wrote the manuscript. FTLM, MJH and CL developed the liver MPS COMSOL Multiphysics model.

## Acknowledgements

This work was supported in part by NIH UH3 TR000487, NIH S10 program under award number 1S10RR026866-01 and NIH P30 DK026743. The authors acknowledge technical assistance with Seahorse data acquisition from Pete Zushin (UC Berkeley) and use of the cell and tissue analysis facility (CTAF, UC Berkeley). The authors also thank Mary West (UC Berkeley) for assistance with flow cytometry and Simone Kurial flow cytometry data analysis. AL had a fellowship from Novartis Foundation for Medical Bological Research (15B086) and from Fondazione Ettore E Valeria Rossi.

