## Supplementary Materials for "Integrated hiPSC-based liver and heart microphysiological systems predict unsafe drug-drug interaction" (from another version of this preprint: version 1, 2020-05-27)

**Table 1. Cisapride $EC_{50}$ values compiled from the literature.**

The estimated therapeutic plasma concentration (ETPC) of Cisapride is 2-5 nM. MEA: multielectrode array. hiPSC-CMs: human induced pluripotent stem cell derived cardiomyocytes.

| Drug | Description of effect | 2D or 3D | Cell type/ Study model | Analysis | Ion Channel | $EC_{50}$ (nM) | Reference |
|---|---|---|---|---|---|---|---|
| Cisapride | 5-HT4 receptor agonist | 3D | hiPSC-CMs | Voltage dye | hERG | 9.6 | Lee-Montiel et al. (2020) |
| Cisapride | 5-HT4 receptor agonist | 2D | CHO, hERG-transfected | Patch clamp | hERG | 36.5 | 2017[1] |
| Cisapride | 5-HT4 receptor agonist | 2D | HEK-293 | Patch clamp | hERG | 21.2 | 2015[2] |
| Cisapride | 5-HT4 receptor agonist | 2D | HEK-293 | Patch clamp | hERG | 10 | 2016[3] |
| Cisapride | 5-HT4 receptor agonist | 2D | hiPSC-CMs | MEA | hERG | 20 | 2003[5], 2013[6], 2017[4] |
| Cisapride | 5-HT4 receptor agonist | 2D | HEK-293 | Patch clamp | hERG | 20-45 | 2003[5] |
| Cisapride | 5-HT4 receptor agonist | 2D | HEK-293 | Patch clamp | hERG | 20 | 2013[6] |
| Cisapride | 5-HT4 receptor agonist | 3D spheroid | hiPSC-CMs | Calcium oscillation | hERG | 202 | 2017[7] |
| Cisapride | 5-HT4 receptor agonist | 2D | hiPSC-CMs | Calcium oscillation | hERG | 11 | 2017[7] |
| Cisapride | 5-HT4 receptor agonist | 2D | hiPSC-CMs | Patch clamp | hERG | 32 | 2013[8] |
| Cisapride | 5-HT4 receptor agonist | 2D | HEK-293 | Patch clamp | hERG | 27 | 2004[9], 2005[10] |
| Cisapride | 5-HT4 receptor agonist | 2D | COS-7 | Patch clamp | hERG | 240 | 2001[11] |
| Cisapride | 5-HT4 receptor agonist | 2D | HEK-293 | Patch clamp | hERG | 6.5 | 1997[12] |
| Cisapride | 5-HT4 receptor agonist | 2D | HEK-293 | Patch clamp | hERG | 14.4 | 2002[13] |
| Cisapride | 5-HT4 receptor agonist | 2D | hiPSC-CMs | Patch clamp | hERG | 32-35 | 2013[14] |
| Cisapride | 5-HT4 receptor agonist | 2D | HEK-293 | Patch clamp | hERG | 27 | 2005[10] |
| Cisapride | 5-HT4 receptor agonist | 2D | HEK-293 | Patch clamp | hERG | 44.5 | 1997[15] |
| Cisapride | 5-HT4 receptor agonist | 2D | CHO-K1 | Patch clamp | hERG | 23.6 | 1999[16] |
| Cisapride | 5-HT4 receptor agonist | 2D | HEK-293 | Patch clamp | hERG | 18 | 2004[17] |
| Cisapride | 5-HT4 receptor agonist | 2D | HEK-293 | Patch clamp | hERG | 30 | 2004[18] |
| Cisapride | 5-HT4 receptor agonist | 2D | HEK-293 | Patch clamp | hERG | 15 | 2004[19] |

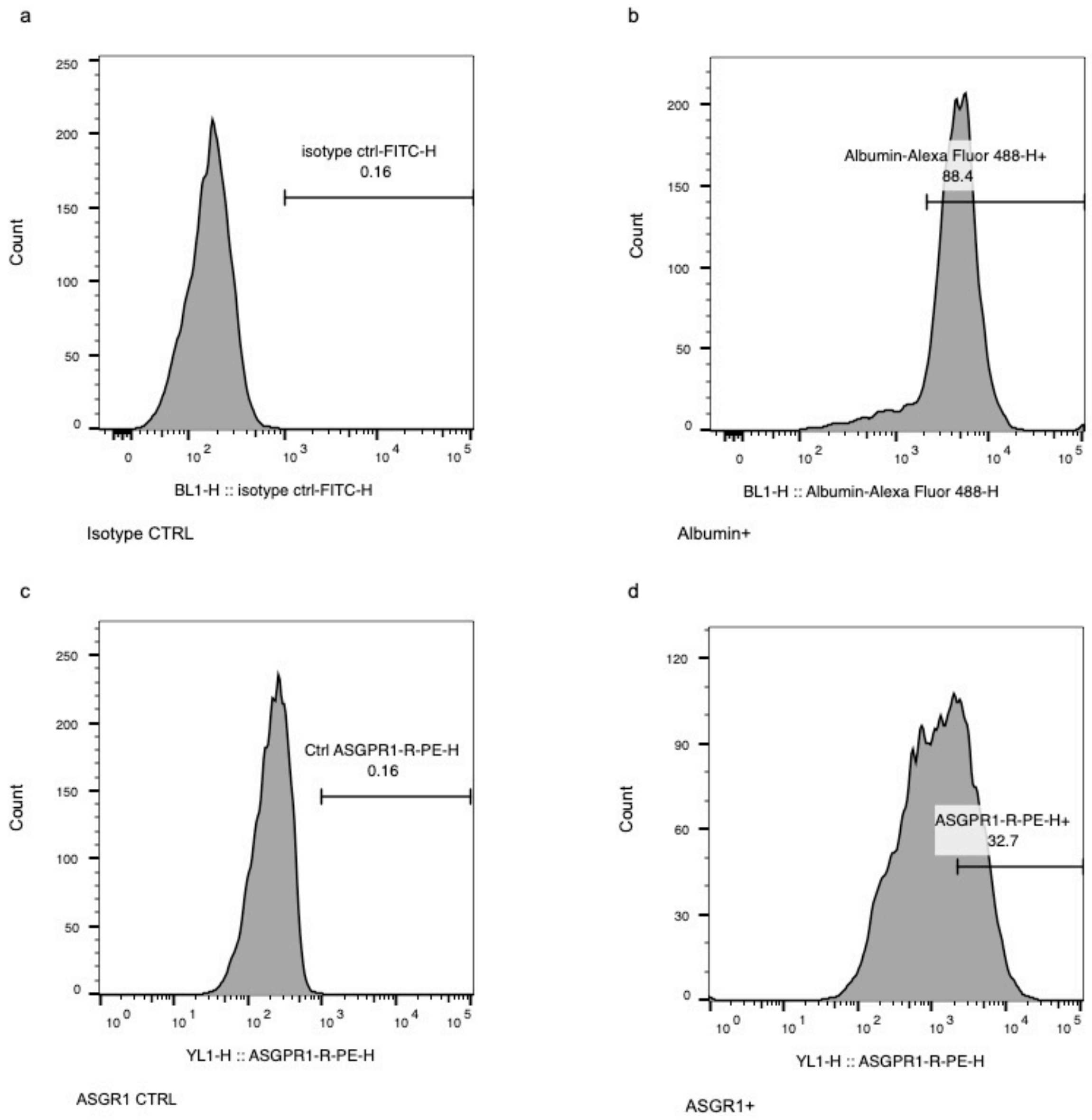


**Supplementary Fig 1. Flow cytometry analysis of hiPSC-Heps.**

Representative flow cytometry results of hiPSC-Heps at day 23 of the differentiation protocol. Isotype control FITC (a), albumin-positive cells (b), ASGR1 control (c) and ASGR1-positive cells (d). Numbers indicate % positive cells.

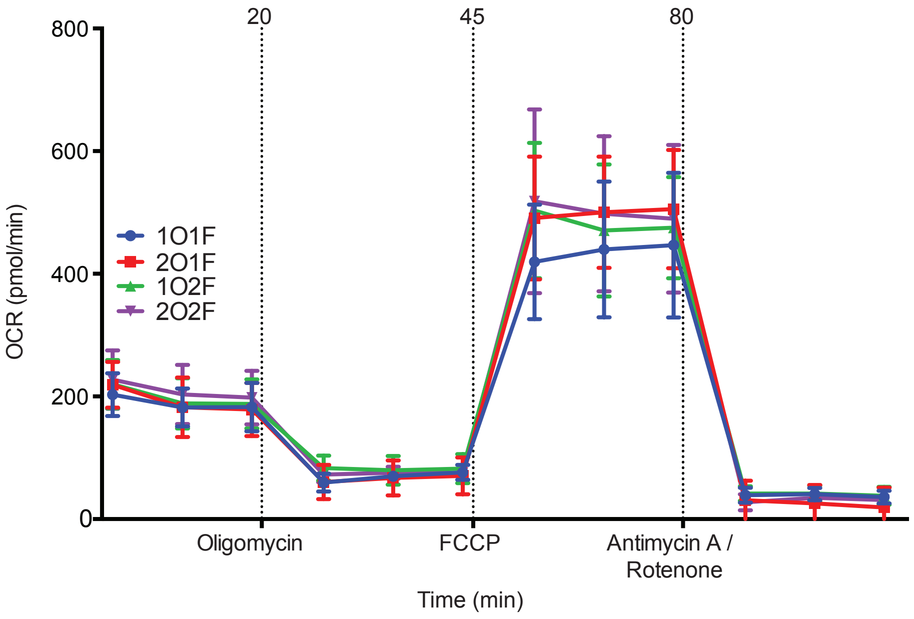


**Supplementary Fig 2. Oxygen consumption rate (OCR) of hiPSC-Heps.**

Direct measurement of OCR was made by sequential exposure of hiPSC-Heps to 1 µM oligomycin (1O1F and 1O2F) or 2 µM oligomycin (2O1F and 2O2F), 1 and 2 µM carbonyl cyanide-4-(trifluoromethoxy)phenylhydrazone (FCCP) and rotenone/antimycin A. Basal respiration was measured for 20 minutes. Oligomycin was injected at 20 minutes to block ATP synthase (complex V), thereby leading to a decrease in electron flow through the electron transfer chain (ETC) and the observed decrease in OCR. FCCP is an uncoupling agent disrupting the mitochondrial membrane potential. FCCP was injected at 50 minutes and its disinhibition of the electron flow through the ETC led to the determined maximal OCR. At 80 minutes a combination of rotenone (complex I inhibitor) and antimycin A (complex III inhibitor) was injected to shut down mitochondrial respiration.

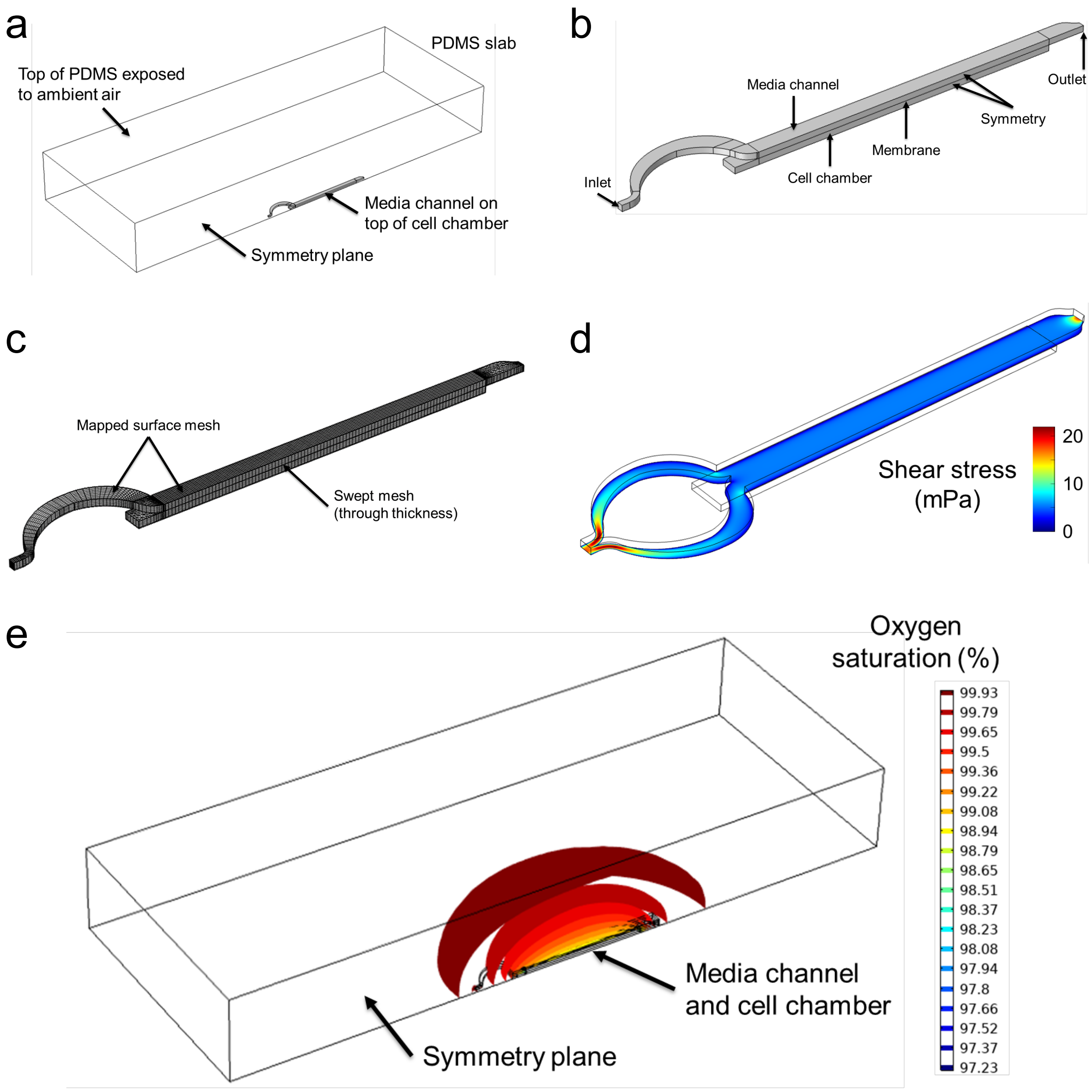


**Supplementary Fig 3. Liver MPS COMSOL Multiphysics® model setup and results.**
Liver MPS COMSOL Multiphysics® model setup (a), liver MPS model setup of cell chamber and media channel (b), liver MPS model mesh of cell chamber and media channel (c), shear stress on membrane (media channel side) at 20 µL/h (d), and oxygen saturation in PDMS at 24 hours (e). The $O_2$ concentrations in the PDMS, and in particular at the media channel and cell chamber walls, are nearly saturated (i.e., above 97% of ambient $O_2$ partial pressure).

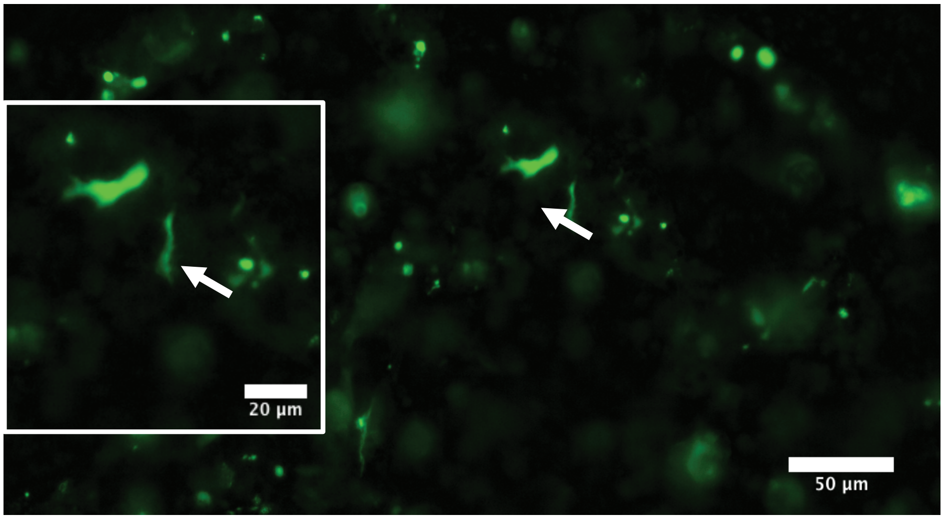


**Supplementary Fig 4. Stable differentiation and function of hiPSC-Heps in liver MPS.**
hiPSC-Heps form bile canaliculi after 2 weeks in culture inside the liver MPS as shown by MRP2-dependent intracanalicular transport and accumulation of the metabolite glutathione methylfluorescein one hour after incubation with 5-chloromethylfluorescein diacetate.

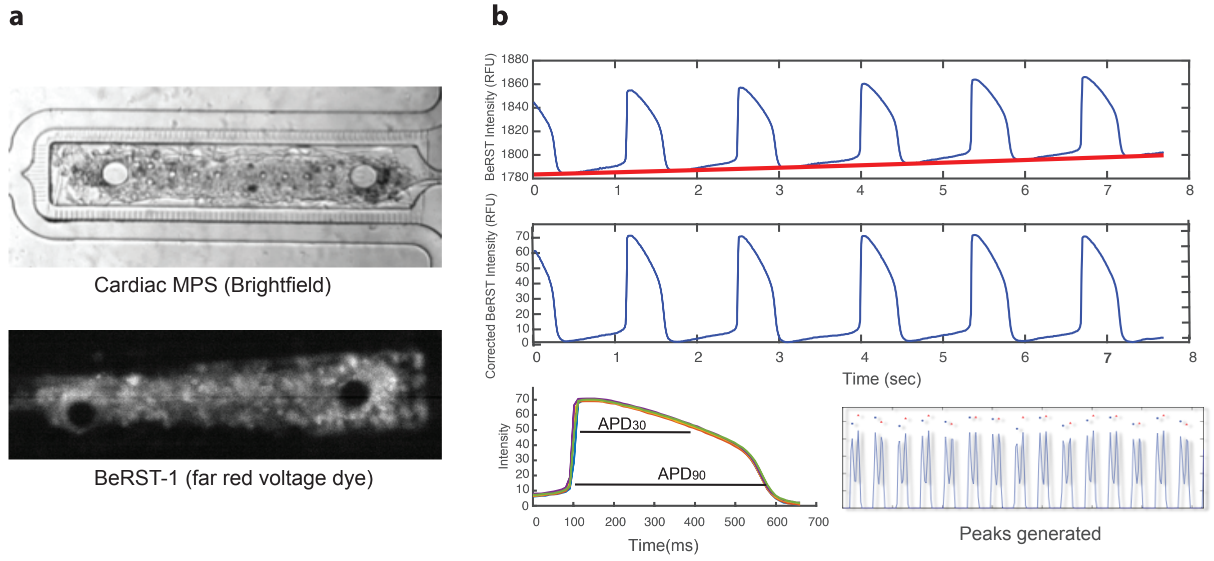


**Supplementary Fig 5. Voltage-sensitive dye BeRST-1 and MATLAB motion tracking analysis software.**

(a) Cardiac MPS brightfield image (top) and labeling with far red voltage dye BeRST-1 (bottom).

(b) Data sample of quantification of the beat rate and APD signal. BeRST-1 videos were analyzed for beat rate and 80% and 90% APD ($APD_{80}$, $APD_{90}$) using custom MATLAB and Python code.

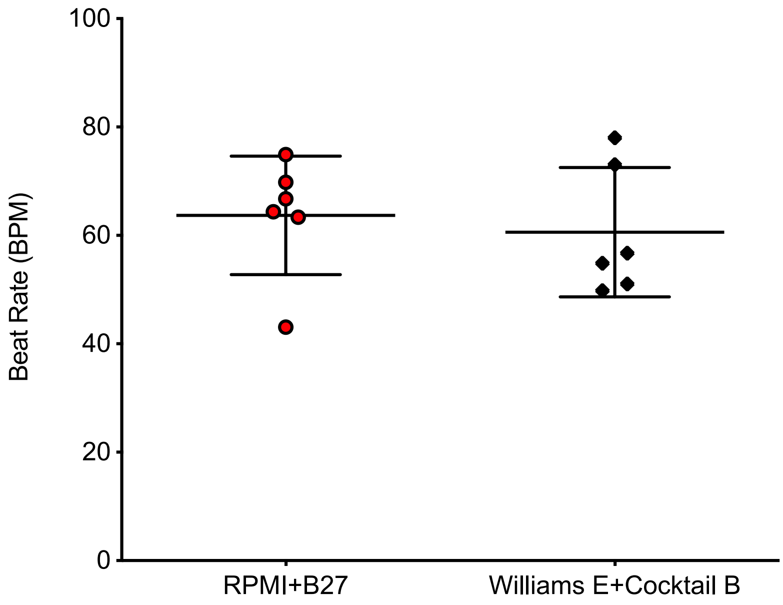


**Supplementary Fig 6. Cardiac effects of common media.**

Effects of different culture media on spontaneous beat rate of hiPSC-CMs in cardiac MPS.

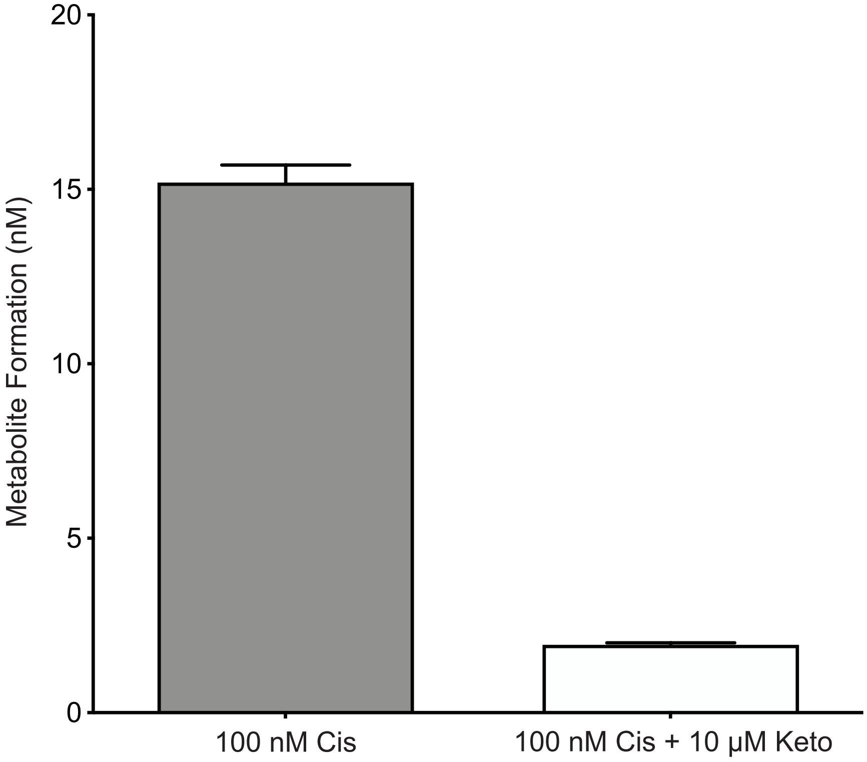


**Supplementary Fig 7. Characterization of low-dose cisapride metabolism in hiPSC-Heps in conventional culture**

Mass spectroscopy analysis of hiPSC-Heps using UFLC-MS/MS shows CYP3A4-mediated cisapride-to-norcisapride metabolism at 100 nM drug input and inhibition by ketoconazole.

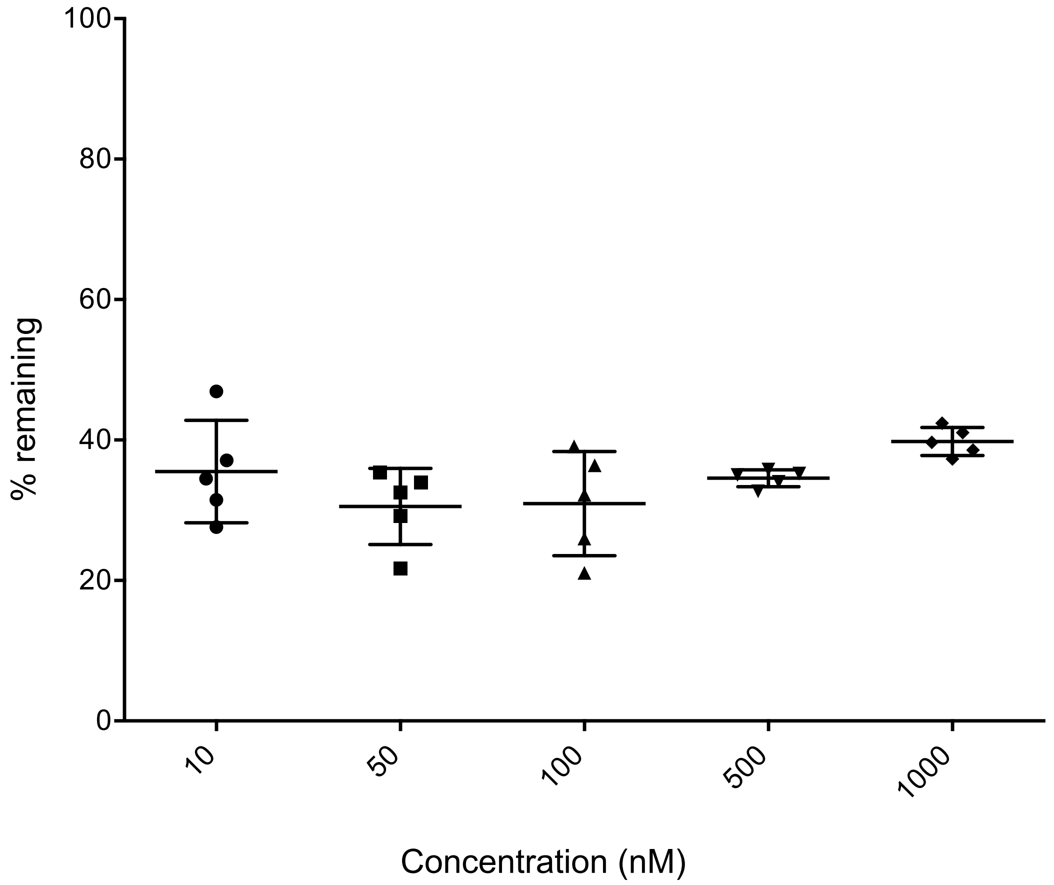


**Supplementary Fig 8. Mass spectrometry analysis of cisapride absorption in PDMS.** Absorption of different concentrations of cisapride in PDMS in the liver MPS was analyzed to correct the concentrations used in the experiments. Ion abundances of the drug in stock samples and in efflux media were measured using LC-MS/MS and the percent remaining was calculated. One device was used per cisapride concentration of 10, 50, 100, 500, and 1,000 nM.

### Supplemental methods

#### Bile Efflux Testing

hiPSC-Heps were tested for normal canalicular efflux of CMFDA at day 7. CMFDA (Life Technologies) was prepared at 4 μM in HCM and added to the liver MPS. The devices were removed from the perfusion pump and imaging was performed using a Nikon Eclipse microscope for collection of 488/525 nm (ex/em) after 1-hour exposure with 30-minute intervals over 1 hour.

#### Oxygen Consumption Rate

Oxygen consumption was analyzed in hiPSC-Heps (on day 25 of differentiation) with the Seahorse XF24 cellular respirometer (Agilent). Mitochondrial stress tests were performed on hiPSC-Heps 100% confluence (80,000 cells/well) in V28 microplates, with XF assay medium supplemented with 1 mM pyruvate (Gibco), 2 mM glutamine (Gibco), and 25 mM glucose (Sigma) at pH 7.4 and sequential additions via injection ports of oligomycin (1 and 2 μM final), FCCP (1 and 2 μM final), and antimycin A/rotenone (1 μM final) during respirometry (concentrated stock solutions solubilized in 100% ethanol [2.5 mM] for mitochondrial stress test compounds). Plates were incubated in a $CO_2$-free incubator at 37°C for 1 hour to allow temperature and pH equilibration.

#### Liver MPS COMSOL Multiphysics® model

The liver MPS COMSOL® model comprising PDMS slab containing media channel and cell chamber separated by a porous PET membrane (**Supplementary Fig. 3a**). By symmetry about the vertical center plane, the computational domain was half the physical domain. The cell chamber base was flush with base of 3.5 mm thick PDMS slab. The top of the PDMS slab was exposed to ambient air with specified oxygen ($O_2$) concentration. The base of device (base of cell chamber and PDMS) was assumed $O_2$ impermeable. $O_2$ flux through PDMS slab sides was also neglected (sides were taken at least 3 slab thicknesses away for this to be a good assumption). The liver MPS model setup of cell chamber, media channel, and membrane portions of computational domain are shown in **Supplementary Fig. 3b**. Media channel and cell chamber heights were 100 μm and the membrane thickness was 15 μm. Flow enters the inlet at 20 μL/h and exits at the outlet. Laminar inflow and outflow conditions were used in COMSOL®. The membrane was modeled as a 2D planar surface between the media channel and cell chamber, with effective diffusivities for $O_2$ and small molecules. Fluid flow across the membrane was neglected, and zero flow velocity was assumed throughout the cell chamber. No-slip conditions were imposed at all walls and membrane surfaces within the medial channel. For $O_2$ transport simulations, cells were assumed to fill the cell chamber volume at a specified density and consume $O_2$ at a specified rate (see main text for details). Liver MPS model mesh of the cell chamber, media channel, and membrane portions of computational domain are shown in **Supplementary Fig. 3c**. Mapped meshes were used to control the axial and lateral mesh resolutions on the straight and curved portions of the media channel. Triangular mesh elements were used at connection points, and surface meshes were then swept downward through the thicknesses of the media channel and cell chamber (5 elements through each). Mesh sensitivity analysis was performed to show that this mesh resolution yields sufficiently accurate numerical results. Quadratic finite elements are used for the flow velocity components and species concentration, while linear elements are used for fluid pressure. For shear stress calculations on the membrane (media channel side) at 20 μL/h, no-slip conditions were assumed at all walls as well as along the membrane (**Supplementary Fig. 3d)**. $O_2$ saturation in PDMS was determined at 24 hours for a flow rate of 20 μL/h. (**Supplementary Fig. 3e**).
